# Convergent evolution of primate testis transcriptomes reflects mating strategy

**DOI:** 10.1101/010553

**Authors:** Etka Yapar, Ekin Saglican, Handan Melike Dönertaş, Ezgi Özkurt, Zheng Yan, Haiyang Hu, Song Guo, Babür Erdem, Rori V. Rohlfs, Philipp Khaitovich, Mehmet Somel

**Affiliations:** Department of Biological Sciences, METU, Ankara, Turkey; European Molecular Biology Laboratory, European Bioinformatics Institute, Wellcome Trust Genome Campus, Hinxton, Cambridge, UK; Quadram Institute Biosciences, Norwich Research Park, Norwich, UK; Earlham Institute, Norwich Research Park, Norwich, UK; CAS Key Laboratory of Computational Biology, CAS-MPG Partner Institute for Computational Biology, Shanghai Institute of Nutrition and Health, Shanghai Institute for Biological Sciences, University of Chinese Academy of Sciences, Chinese Academy of Sciences, 320 Yue Yang Road, Shanghai, 200031, China; Department of Biology, San Francisco State University, San Francisco, California, USA; Center for Neurobiology and Brain Restoration, Skolkovo Institute of Science and Technology, Moscow, Russia

## Abstract

In independent mammalian lineages where females mate with multiple males (multi-male mating strategies), males have evolved larger testicles relative to those lineages where females mate with fewer males (single-male mating strategies). Here we study published bulk testis transcriptomes from humans, chimpanzees, gorillas and rhesus macaques, as well as mice and rats. Employing a formal model of adaptive evolution, we find that testis transcriptomes have also evolved convergently, reflecting each species’ mating strategy. Using deconvolution, we infer that testis transcriptome divergence patterns largely reflect convergent shifts in tissue cell type composition. However, we also identify modest amounts of convergent evolution at the cell-autonomous level by analyzing cell-type specific transcriptome data from spermatids and spermatocytes. We further show that in the single-male mating primates, human and gorilla, testis transcriptome profiles are paedomorphic relative to those of multi-male primates, chimpanzee and macaque, suggesting that shifts in timing or rate of testis development could underlie convergent changes in testis mass, histology, and transcriptomes.

## INTRODUCTION

Molecular investigations of hominid evolution frequently focus on the human brain, which has tripled in size since our last common ancestor with chimpanzees (Carroll, 2003; Sherwood et al., 2012; Herculano-Houzel, 2012). Despite this pre-occupation with increasing brain size on the hominid lineage, a more striking morphological change is in decreasing testes size. Chimpanzee testicles (150-170 g) can be 3-10 times larger than human testicles (16-50 g) (Schultz, 1938; Dixson, 2012; Harcourt et al., 1981), a shift is even more striking when considering that humans are 30% larger than chimpanzees in body size.

It has been hypothesized that interspecies testis mass differences could be explained by their mating strategies. In multi-male or promiscuous mating strategies, where females regularly mate with multiple males during an estrous cycle (e.g. polyandry or polygynandry), inter-male sperm competition is prevalent, and is expected to select for higher copulation frequency, higher ejaculate volume, and larger testis mass (Short, 1979). This could explain large testicles in the chimpanzee, with their well-documented multi-male mating patterns, as compared to humans or gorillas, which display “single-male” mating patterns, where female mating with multiple males during a single estrous cycle is less common (e.g. polygyny or monogamy), (Dixson, 2012). Testing this notion four decades ago, Harcourt and colleagues found a marked relationship between relative testis mass (residual testis weight accounting for body weight) and mating systems across primates, such that multi-male species had higher relative testis mass than single-male species (Harcourt et al., 1981). Later work corroborated this relationship between relative testis mass and mating strategy across diverse mammalian taxa (Dixson, 2012; Kenagy and Trombulak, 1986; Ramm et al., 2005; Hosken, 1997; Harcourt et al., 1995).

The distribution of mating systems and relative testis mass within the primate phylogeny suggests that large and small testicles evolved multiple times, convergently. Testis histology and spermatogenesis rates may also be evolving convergently. Testicles of multi-male primates contain 1.5-3 times higher proportions of seminiferous tubules to interstitial (connective) tissue, compared to ratios close to 1:1 in single-male primates (Schultz, 1938; Harcourt et al., 1981). Species with large relative testis mass also have higher rates of spermatogenesis than other species (Ramm and Stockley, 2010). These observations suggest that the evolution of testis anatomy and histology can be rapid, possibly more so than in other tissues. Strong positive selection on reproductive phenotypes (Dixson, 2012; Short, 1979; Stockley, 2004) and relatively modular organization of testis development (Stockley, 2004; Sekido and Lovell-Badge, 2013) could be reasons behind rapid evolution of testis phenotypes.

Testis evolution also reveals unique patterns at the transcriptome level. Comparing humans and chimpanzees using microarrays, Khaitovich and colleagues found significantly higher gene expression divergence and low within-species diversity in the testis relative to other tissues, including the brain and the liver (Khaitovich et al., 2005, 2006b). Furthermore, protein coding sequences of testis-expressed genes were found to evolve faster than those of other genes (Khaitovich et al., 2005, 2006a). Comparing mammalian tissues using RNA-sequencing (RNA-Seq), Brawand and colleagues further highlighted the testis as the most rapidly evolving among six tissues at the expression level (Brawand et al., 2011). They also noted an apparent similarity between human and gorilla testis transcriptomes (two single-male species) relative to the chimpanzee, although this effect was not quantified. Meanwhile, a recent study on gene expression changes across development in five tissues across different mammals (Cardoso-Moreira, 2019) reported testis as the tissue with the highest number of genes with newly acquired developmental gene expression patterns (in rodents and the rabbit, though not in humans).

Here we study the evolution of adult testis transcriptomes in the context of mating behavior differences among four closely related primate species: human, chimpanzee, gorilla and macaque. We reanalyse published transcriptome data sets employing a formal model of adaptive phenotypic evolution, and test whether bulk testis transcriptome profiles have evolved in convergent fashion among species with similar mating strategies. We next use cell type-specific transcriptome data and deconvolution analysis to investigate whether bulk tissue level expression divergence between species is predominantly caused by cell-autonomous regulatory changes, or by changes in tissue composition. Finally, we investigate possible effects of developmental timing and rate differences on tissue composition change and expression divergence.

## RESULTS AND DISCUSSION

### Testis transcriptomes mirror species’ mating strategies

We hypothesized that single-vs. multi-male mating strategies, which evolved in distinct primate lineages (Figure 1A), and which supposedly led to convergent evolution of relative testis mass, could also be reflected in bulk testis transcriptome profiles. If so, we would expect convergent patterns of gene expression evolution, which might explain aforementioned observations on testis expression divergence in primates. For this, we combined adult testis bulk tissue transcriptome data from two studies (Khaitovich et al., 2005; Brawand et al., 2011). These data sets together comprise 8 humans, 7 *Pan* genus individuals (6 common chimpanzees and one bonobo, that we together refer to as “chimpanzees”), one gorilla, and two rhesus macaques; all Old World anthropoids. Here, the human and gorilla represent historically single-male species, as testified by their low relative testis mass, whereas chimpanzee and rhesus macaque represent multi-male species (Harcourt et al., 1981; Dixson, 2012). This combined primate testis transcriptome dataset contained *n*=7305 one-to-one orthologous primate genes expressed in the testis.

**Figure 1:**
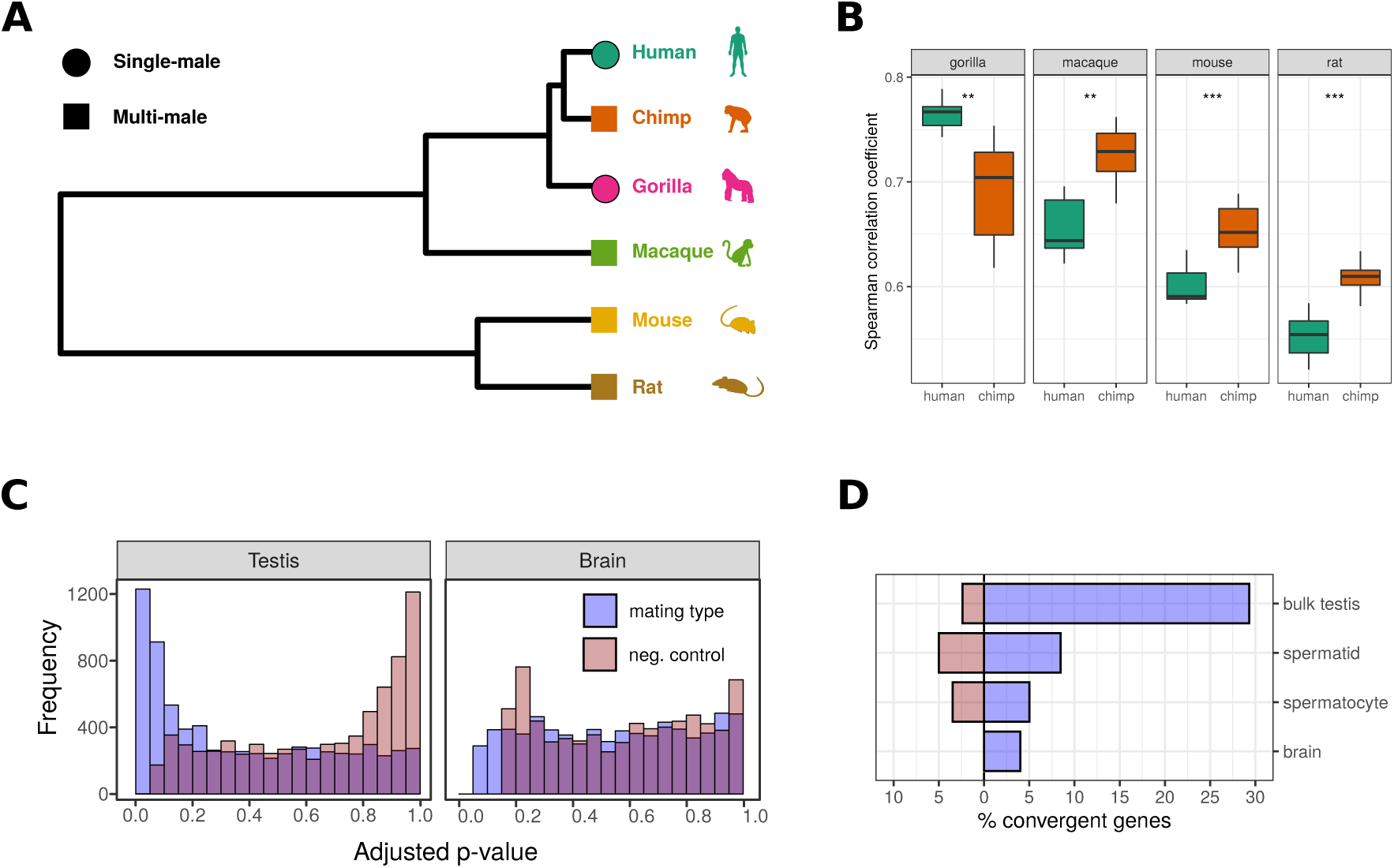
**(A)** Phylogenetic tree of species included in this study. **(B)** Overall correlations of gorilla, macaque, mouse and rat to human versus chimpanzee testis transcriptome profiles. The correlations were calculated across differentially expressed genes between human versus chimpanzee; n=4295 in analyses involving gorilla or macaque, and n=2847 in analyses involving mouse or rat. For each species tested, we tested for a significant difference between correlations to human versus chimpanzee using a two-sided permutation test based on the Mann-Whitney U statistic [**(**):** *p <* 0.005, **(***):** *p <* 0.001]. **(C)** Adjusted p-value distributions of convergent evolution tests with EVE for the testis and brain. Blue shaded histograms show the p-value distribution for the tests of mating strategy-related convergent evolution [(human-gorilla) vs. (chimp-macaque)], whereas the red shaded histograms show those of negative controls [(human-macaque) vs. (chimpgorilla)]. **(D)** Percentage of convergent genes among the bulk testis, spermatocyte, spermatid and the brain datasets.

We used two approaches to investigate convergent evolution patterns. In the first approach, we treated human and chimpanzee as representatives for single-vs. multimale mating, respectively, and compared gorilla and macaque to this pair. Under the null hypothesis of no mating strategy effect, given the phylogeny (Figure 1A), we would expect both gorilla and macaque to be equally similar to humans and chimpanzees. If a mating strategy effect exists, gorilla should be closer to humans, and macaques to chimpanzees. To test this, we focused on *n*=4295 orthologous genes differentially expressed between human and chimpanzee (two-sided Mann-Whitney U test, Benjamini-Hochberg (BH) corrected q<0.10). Using Spearman correlation, we then compared the gorilla and rhesus macaque testis transcriptome profiles with those of humans and of chimpanzees across these differentially expressed genes (Figure 1B). The gorilla testis transcriptome showed significantly higher correlation with those of humans than of chimpanzees; the reverse was true for the macaque (permutation test, p<0.04), as would be expected based on their mating strategy affinities.

We then extended this correlation analysis to include additional species. We first included mouse (*Mus musculus*) and rat (*Rattus norvegicus*); both species display well-documented multi-male mating behavior (Kenagy and Trombulak, 1986; Chalmel et al., 2007). Consistently, their testis transcriptomes also showed higher correlation with those of chimpanzees than of humans (Figure 1B; permutation test, p<0.001) (note that mouse and rat are related lineages and therefore do not represent fully independent comparisons). We further included published testis transcriptome data from the common marmoset (*Callithrix jacchus*), gray short-tailed opossum (*Monodelphis domestica*) and platypus (*Ornithorhynchus anatinus*). We found that the marmoset and opossum testis samples showed no preference towards either human or chimpanzee, while the platypus showed higher affinity toward the human (Figure S3). Due to ambiguities with regard to the sampled individuals’ characteristics (age and breeding status) and/or the species’ mating behavior, however, we decided not to include these latter three species in downstream analyses (see Methods for discussion).

### An Ornstein-Uhlenbeck model-based analysis identifies common convergent evolution in testis transcriptomes

We next investigated convergent evolution of testis expression levels using a formal statistical framework, the Expression Variance and Evolution (EVE) model (Rohlfs and Nielsen, 2015). EVE relies on the Ornstein-Uhlenbeck model of phenotypic evolution, while also taking into account intraspecies variance. EVE allows us to calculate the likelihood of expression levels under (a) a null model of uniform stabilizing selection towards a common expression level across all lineages, and (b) an alternative model convergent evolution towards varying expression levels in *a priori* determined lineages.

We focused on the four Old World anthropoids. Given their phylogenetic relationships (Figure 1A), gene expression levels that systematically group chimpanzees with rhesus macaques, and humans with gorillas, represent convergent patterns that could be driven by the species’ mating system differences, and could be detected by EVE. Testing each of the *n*=7,305 genes in the combined primate testis transcriptome dataset, 2,143 (29.3%) displayed significant convergent patterns with respect to mating strategy (chi-square test, at BH corrected q<0.10; Figure 1C; Table S2). As negative control, we also tested for convergent patterns that cluster macaques with gorilla, and chimpanzees with humans (Figure 1C). This yielded only 175 genes (2.4%) at q<0.10.

If the observed convergent evolution signatures are indeed related to mating strategy evolution, these signatures should be prevalent in the testis but not in other tissues. We therefore tested whether convergent mating strategy-related expression patterns may be as widespread in the brain. We performed the same analysis using prefrontal cortex transcriptome data comprising the same four species, using a similar but larger sample than the testis data (Methods). Out of *n*=7,212 orthologous genes only 289 (4%) showed significant convergence with respect to mating strategy (Figure 1C). This result supports the notion that convergent evolution of primate testis transcriptomes is related to their convergent mating strategy characteristics.

### Widespread convergent evolution of testis cell type composition

How might we explain parallel expression changes across thousands of genes in distinct lineages? One simple scenario could involve parallel shifts in tissue cell type composition, i.e. tissue architecture. Indeed, compared to humans and gorillas, chimpanzee testicles contain a higher concentration of seminiferous tubules than interstitial tissue, a feature that possibly correlates with the chimpanzee’s higher rate of sperm production (Dixson, 2012; Harcourt et al., 1981; Stockley, 2004; Fujii-Hanamoto et al., 2011). Similar differences have been noted among *Mus* species with varying levels of female promiscuity at the histological level (Montoto et al., 2012), which appear also reflected at the bulk transcriptome level (Good et al. 2010).

During germ cell maturation, there occurs a switch in global expression at meiosis I. This creates pronounced transcriptomic differences between (a) meiotic/post-meiotic germ cells, which we here denote POST, and which include spermatocytes and spermatids, and (b) pre-meiotic/somatic cells, which we here denote PRE, and which include Sertoli cells, spermatogonia, and interstitial cells in the testis (Figure S5) (Chalmel et al., 2007). The seminiferous tubule transcriptome mainly reflects POST cell expression in rodents and in humans (Schultz et al., 2003; Chalmel et al., 2007). Thus, in multi-male species’ testicles, where we expect higher seminiferous tubule:interstitia ratios (Dixson, 2012; Harcourt et al., 1981), we also expect higher POST:PRE cell type ratios per unit volume. We hypothesized that POST vs. PRE cell type shifts may have evolved convergently in primates, and that such shifts could explain the observed convergent expression evolution patterns in bulk testis transcriptomes.

We tested this hypothesis employing deconvolution analysis based on linear regression. We used conserved PRE and POST transcriptome profiles derived from mice and we predicted the relative proportions of PRE and POST cells within bulk testis samples of the primate and rodent species described earlier (Methods). POST expression was exclusively dominant in multi-male, whereas PRE expression was exclusively dominant in single-male species’ testis transcriptomes (Figure 2A). Further, using the EVE method, we found significant evidence for convergent evolution of predicted POST/PRE ratios under single-vs. multi-male mating systems (p=0.0006 for primates, p=0.0008 for primates and rodents combined; Methods).

**Figure 2:**
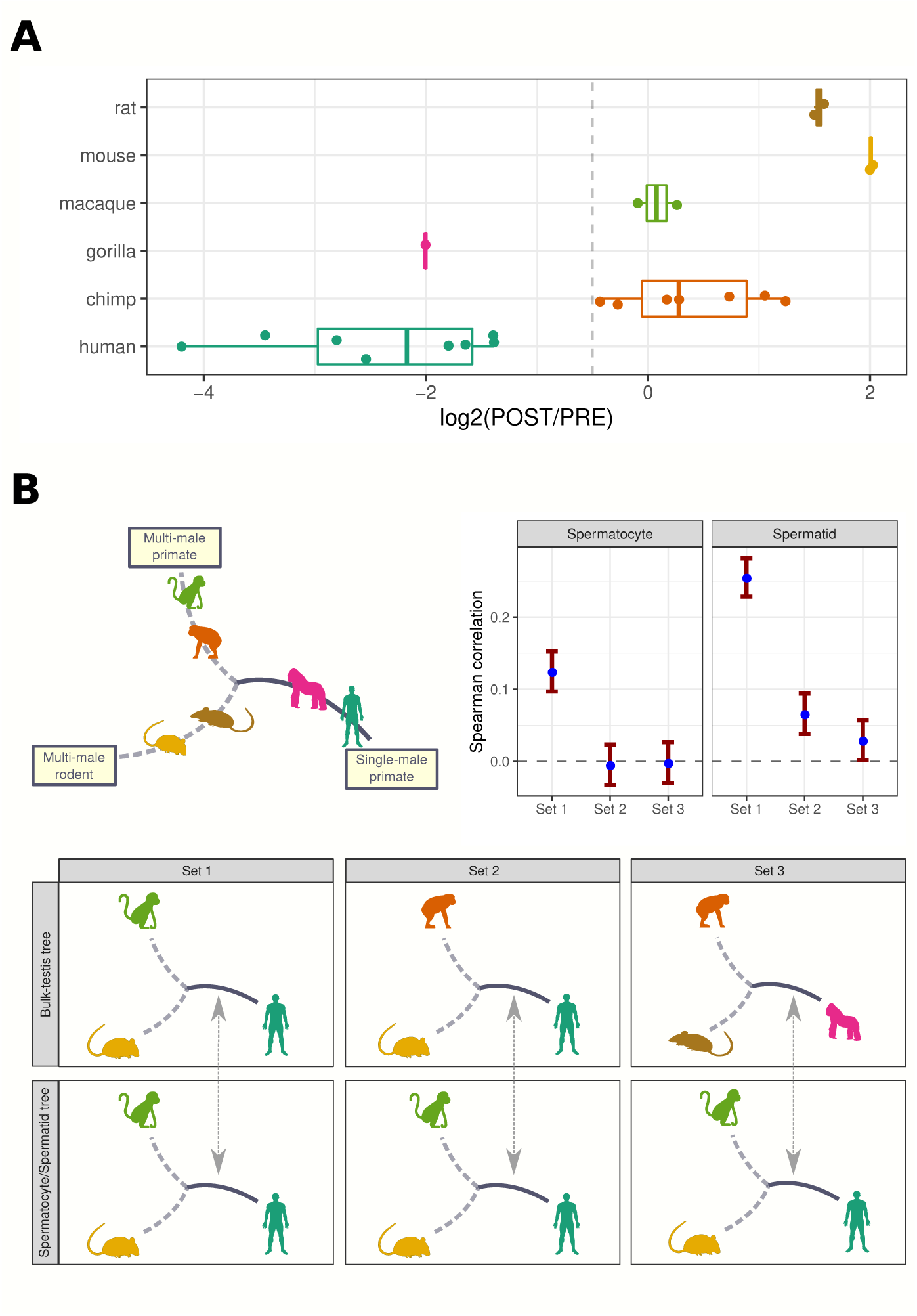
**(A)** Results of cell-type deconvolution analyses of adult bulk testis samples. Boxplots show the distribution of *log*_2_(*b_POST_/b_PRE_*) ratios across individuals of each species. **(B)** Results of branch length-based comparative analysis of cell-autonomous expression changes and bulk testis expression changes. For each gene present in either the spermatocyte/spermatid or bulk testis datasets, a neighbor-joining tree was built based on gene expression levels of the indicated species. In each case, the length of the single male primate branch (either human or gorilla) was calculated. The single-male branch length distributions calculated from bulk testis data and from spermatocyte/spermatid data were then compared using Spearman correlation across 4894 genes, for Sets 1, 2 and 3, and for spermatocytes and spermatids, separately. The chart on the top-right shows the Spearman correlation coefficient estimates between the single-male branch lengths of two datasets across different tree sets used, shown on the x-axis. Ranges indicate the 95% confidence intervals calculated by bootstrapping genes.

We performed two additional analyses to gauge the effect of tissue composition shifts on the observed convergent transcriptome changes. In the first, we calculated (a) effect size between single-vs. multi-male primate species on primate bulk testis gene expression, for each gene, and (b) PRE vs. POST cell type effect size on expression of the same genes (Methods). The two effect size measures were correlated (Spearman’s rho=0.40, p<10^-15^; Figure S6A). Second, we found that genes displaying significant convergent evolution in primate testis transcriptome data also have higher PRE vs. POST cell type absolute effect sizes, relative to genes showing no indication of convergent evolution (Mann-Whitney U test p<10^-7^; Figure S6B).

These results demonstrate that evolutionary shifts in tissue composition significantly contribute to the mating strategy-related convergent expression patterns observed in bulk testis transcriptomes. An additional contributing source could be cell-autonomous expression changes that might also evolve in convergent fashion, as we explore in the following.

### Convergent evolution signals at the cell-autonomous level

To investigate convergent evolution within specific cell type transcriptomes, we took advantage of a published dataset of purified pachytene spermatocyte and round spermatid expression profiles from adult humans, rhesus macaques and mice (Lesch et al. 2015); this dataset we will henceforth refer to as the ‘cell type-specific’ dataset. We first used EVE to test each gene for convergent expression patterns in the macaque-mouse pair, i.e. genes for which these two multi-male species have more similar expression levels than expected given the phylogeny of the three taxa. In the spermatocyte dataset we found 5% and in the spermatid dataset 8.5% of genes with signs of convergent evolution according to mating strategy (chi-square test, at BH corrected q<0.10). We then ran EVE models of human-mouse convergent expression as negative control, which yielded 3.5% and 5% of genes in the spermatocyte and spermatid datasets, respectively. The transcriptome-wide likelihood ratio distributions for the mating strategy-related convergence model were significantly higher than those under the negative control model, both for spermatocytes and spermatids (one-sided Wilcoxon signed-rank test, p<10^-10^, Table S1).

Although the proportion of convergently evolving genes (5% and 8.5%) in spermatocytes and spermatids is modest compared to convergently evolving genes in bulk testis transcriptomes using the four Old World Anthropoids (29.5% of genes), this difference could be partly due to the small size of the cell type-specific datasets, which include only three and more distantly-related species. Indeed, when we implemented EVE on bulk testis transcriptomes from human, rhesus macaque and mouse (Methods), we found 14.5% of genes showing mating strategy-related convergence signatures, while 5% were significant in the negative control (Table S1). These results suggest that cell-autonomous expression may also show mating strategy-related convergent evolution.

If spermatocyte or spermatid transcriptomes evolve convergently driven by mating strategy, then human vs. macaque cell autonomous expression divergence may be correlated not only with bulk testis expression divergence between human vs. macaque, but also with bulk testis expression divergence between other single-vs. multi-male species pairs (e.g. gorilla vs. chimpanzee). We tested this idea using two approaches.

First, we built neighbor-joining trees of gene expression for each gene, using both the bulk testis and cell type-specific datasets. Each tree includes one single-male primate, one multi-male primate and one multi-male rodent, using available species combinations per dataset (Figure 2B). We found that, using the spermatid data, single male branch lengths consistently display positive correlation with their bulk testis counterparts irrespective of the species trio tested (bootstrap 95% confidence interval > 0; Figure 2B). Hence, the correlations appear to be driven by mating strategy rather than only species identity, and are thus consistent with mating strategy-related convergent evolution of spermatid expression. Interestingly, this was not observed for spermatocytes.

In a second approach, we used multiple regression models where we jointly studied the effects of tissue composition shifts and cell-autonomous changes in bulk testis data. We built three multiple linear regression models in which the response variables were (i) human vs. macaque, (ii) human vs. chimpanzee, or (iii) gorilla vs. chimpanzee bulk testis effect sizes, across all expressed genes. Note that species pairs were chosen to be single-vs. multi-male. We included three explanatory variables in each model: (a) human vs. macaque effect sizes in spermatocytes, (b) human vs. macaque effect sizes in spermatids, (c) PRE vs. POST effect sizes, across the same genes (Methods). We found that PRE vs. POST effect size, or tissue composition, was a reliable estimator in all three models (p<10^−82^ for all three models, Table S2). In model (i), human vs. macaque effect sizes in spermatocytes and spermatids were also significant (p<10^−7^), although with smaller slope compared to PRE vs. POST effect sizes. Spermatid effect size was also significant in model (ii) but not model (iii), while spermatocyte effect size was significant in neither model (ii) nor (iii) (Table S2). For instance, human vs. macaque effect size in spermatids was a relevant estimator for human vs. chimpanzee bulk testis effect size, although not for gorilla vs. chimpanzee bulk testis effect size. The differences between the branch length correlation results and the regression results could be related to the use of a multi-male outgroup in the former analysis, thereby increasing statistical power.

We noticed that correlations between bulk testis and cell type-specific transcriptome effect sizes are stronger using spermatid expression profiles, as opposed to spermatocyte profiles. This may be partly caused by spermatid cells being more abundant than spermatocytes within seminiferous tubules, at least in humans (Skakkebaek and Heller, 1973). Spermatids also show consistent positive correlation with the bulk testis data in the branch length-based analysis. In addition, EVE results for spermatids reveal a sizeable fraction of convergently evolving genes, more than those in spermatocytes. The data thus suggest that cell-autonomous expression changes may be evolving convergently driven by mating strategy, at least in spermatids. Meanwhile, the magnitude of cell-autonomous convergent expression appears considerably smaller than convergent expression changes attributable to tissue composition shifts.

### Paedomorphism in testis transcriptomes of single-male species

In a lineage that evolved a multi-male mating system, males could evolve larger testicles by extending or accelerating progenitor germ cell divisions during adolescence, thus increasing both the number and proportion of germ-line cells in the testis and amplifying sperm production (Montoto et al., 2012). Alternatively, in lineages that evolve single-male mating, genetic variants that slow down germ cell proliferation may fix by drift or by selection, leading to a paedomorphic state relative to those of multi-male species. Note that here we cannot estimate the ancestral state and therefore cannot determine whether hypermorphosis or paedomorphism could be causal for the observed differences (Rice 1997); we rather refer to paedomorphism as any character state that resembles the juvenile state of related species.

We hypothesized that shifts in the rate or timing of development should also manifest themselves also at the transcriptome level. To test this, we first used a published testis development transcriptome series, where bulk testis tissue had been sampled from newborn to adult mice (Schultz et al., 2003). We compared this dataset with transcriptome profiles from the adult human, chimpanzee, gorilla and macaque testes, as well as PRE and POST cell types (Methods). For each species or cell type, we determined the age of mice it showed highest overall affinity to, using Spearman correlation. As expected, PRE cells showed highest correlation to pre-adolescent mice, and POST cells, to adult mice (Figure 3A). We further found that adult chimpanzee and macaque testes show highest affinity to those of adult mice (median=41 days), while human and gorilla adult testes show highest affinity to subadult mice (median=23 days, Figure 3A). We confirmed that timing differences between humans and chimpanzees data was significant (permutation test, p<10^-5^; Methods). Adding macaque and gorilla to this comparison did not change the result (permutation test, p<10^-5^).

**Figure 3:**
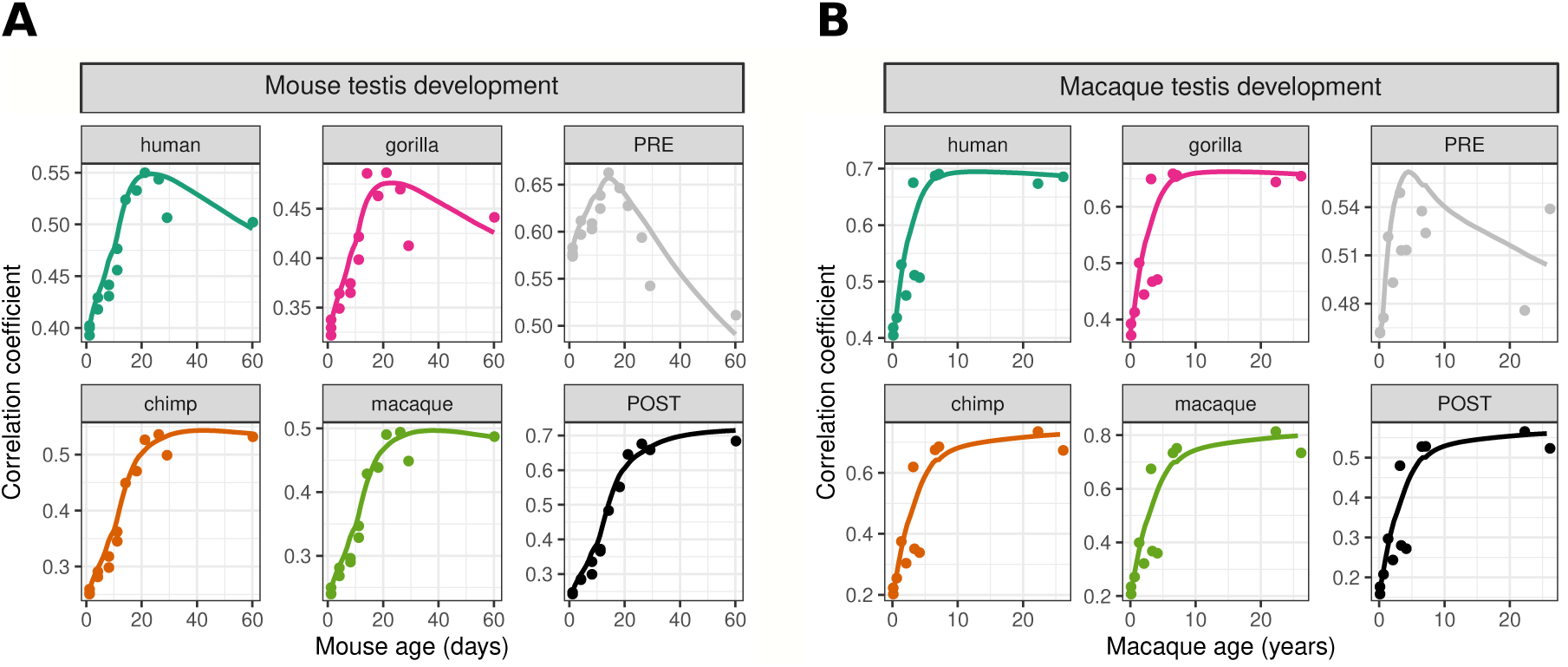
Correlations between adult primate bulk testis expression profiles and PRE and POST cell-type profiles to **(A)** mouse and **(B)** macaque testis developmental profiles. The correlations were calculated across 2295 and 2321 genes with significant Spearman correlation with age in mouse and macaque, respectively. Across mouse testis development, human and gorilla show the maximum correlation to juvenile/adolescent mice (median=23 days of age), whereas chimpanzee and macaque show the maximum correlation to adult mice (median=41 days). There is a trend in the same direction when the same species are compared across the macaque testis development (median=14 years of age and median=25 years of age, respectively). For both cases, differences between the age of maximum correlation for single- and multi-male species (as well as human and chimp differences), are statistically significant (permutation test using Mann-Whitney U statistic, *p <* 10*^−^*^5^ for all comparisons).

To investigate this trend further, we generated microarray data from bulk testis samples from 12 macaques ranging from newborns to old adults (Methods). Performing the same analysis as above, we found a trend in the same direction (Figure 3B; p<10 ^-5^): both datasets support the notion that human and gorilla testis transcriptome profiles are paedomorphic relative to those of multi-male species.

Previous work on comparative brain development found that humans display neotenic expression patterns relative to chimpanzee and macaque in neuron-related gene sets (Somel et al., 2009; Liu et al., 2012). We therefore asked whether the observed transcriptome-wide paedomorphism in human and gorilla testes could be also observed in the brain. To test this, we generated a mouse neocortex development transcriptome dataset, including 8 mice with ages ranging from newborns to 122 days (Methods). We then compared adult human, chimpanzee, gorilla and macaque prefrontal cortex expression profiles to the mouse neocortex development dataset, as above (Figure S8, Methods). In the brain, all adult primates showed highest affinity to young adult mice (∼40 days; Figure S8). In contrast to the testis results, we found no evidence for an overall delay in brain maturation rates between human and chimpanzee (p=0.89). This is in line with the earlier observations that human neoteny in the cerebral cortex comprises specific functional processes, rather than being transcriptome-wide (Somel et al., 2014). Transcriptome-wide paedomorphism in humans and gorillas therefore appears particular to the testicles.

### Molecular regulators of convergent expression patterns

The convergent patterns of testis development and cell type shifts we infer here could be driven by a small number of evolutionary changes in key developmental regulators and their target genes. To investigate this, we studied co-expressed gene groups, sorting genes into six k-means clusters based on their expression profiles across testis development of human, rhesus macaque, and mouse (Figure 4). Two of these clusters showed significant enrichment in convergent genes identified in EVE analysis (Fisher’s exact test, q<0.005; Figure S9). One of them, Cluster #1, was particularly notable in that it showed increasing expression patterns throughout testis development for all three species, and was also enriched in multiple Gene Ontology (GO) Biological Process categories related to spermatogenesis (Table S4; Fisher’s exact test, q<0.1).

**Figure 4:**
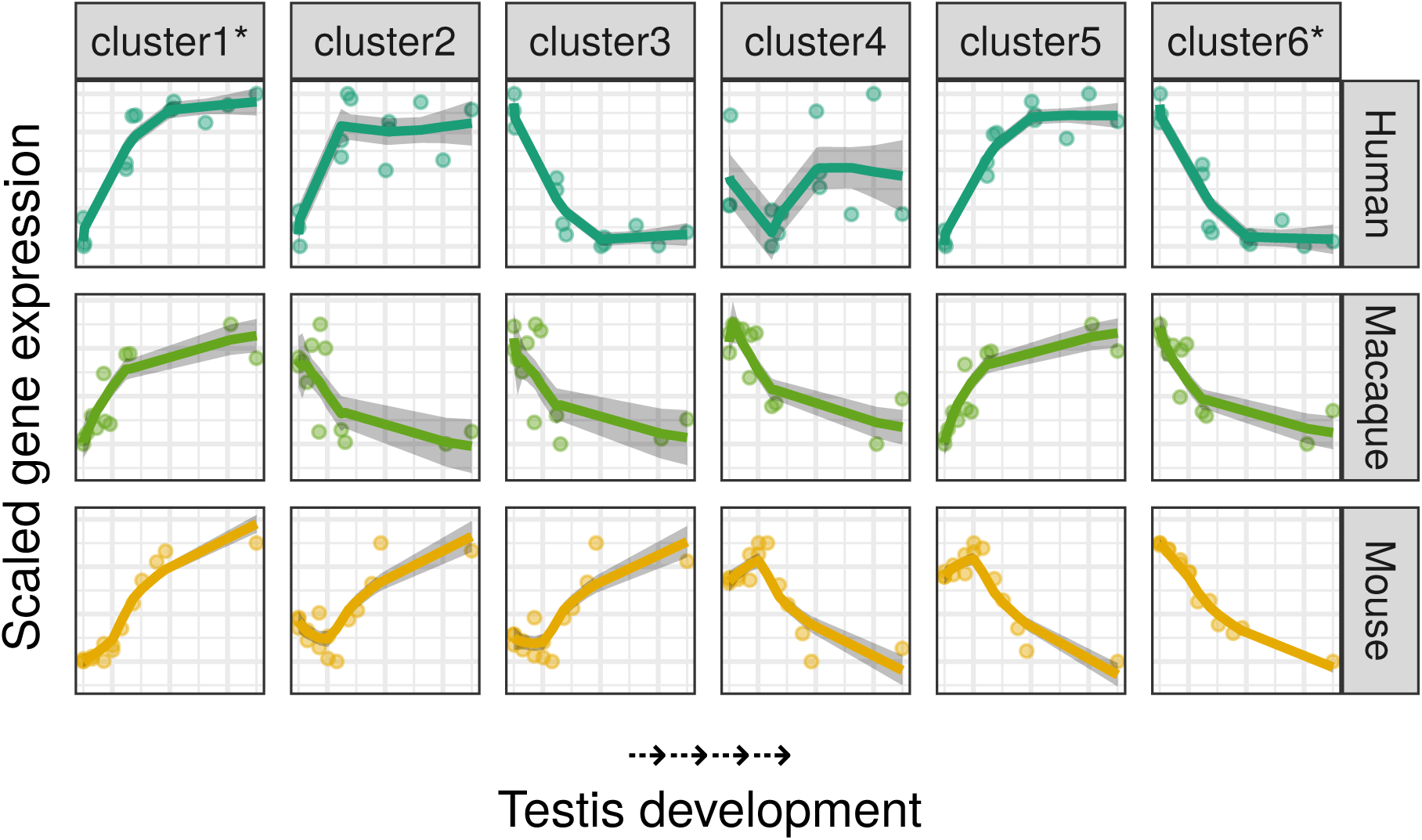
Patterns of expression change throughout human, macaque and mouse testis development. The y-axes show mean expression levels of genes in 6 k-means clusters. The expression levels are scaled to mean 0 and standard deviation 1 before calculating the means. The x-axis shows age, from newborns/infants to adults of each species. Clusters marked with an asterisk are enriched in convergent genes (Fisher’s exact test, *q <* 0.10). Human samples span ages from 7 months to 55 years, macaque samples span ages 16 days to 26 years, mouse samples span ages 1 day to 2 months. Six k-means clusters include 960, 720, 1120, 790, 688 and 1318 genes, respectively.

To determine if a common set of transcription factors regulate the expression of genes in Cluster #1, we performed transcription factor binding site enrichment analysis, using candidate promoter regions estimated based on the human genome. One-hundred-forty candidate transcription factors were enriched in this cluster, out of 475 tested (Fisher’s exact test, q<0.1). Thirty-two of these (22.9%) displayed significant convergent expression patterns in the EVE runs using the adult primate testis dataset (Figure S10). Examining these 32 transcription factors, we noticed three with reported roles in controlling regulation of organ growth: *TEAD1*, *MAX,* and *MXI1*. *TEAD1* is reported as the ultimate target of the Hippo signaling pathway, which is the main evolutionary conserved mechanism for fine-tuning the balance between proliferation and differentiation, and in turn controlling organ growth (Yu and Guan, 2013; Watt et al., 2017). TEAD1 is reported to form a complex with MAX, in a way that each acts as each other’s co-activator (Lin et al., 2017). MAX further forms complexes with MYC(c-Myc) to promote growth and proliferation (Nair and Burley, 2003). Interestingly, MXI1 also forms a complex with MAX in a different setting where it competitively binds to MAX so that there are fewer MAX/Myc complexes to stimulate downstream proliferative pathways (Zervos et al., 1993; Schreiber-Agus et al., 1998). The convergent gene expression patterns found in our adult primate bulk testis data for *MAX* and *TEAD1* seem strange at first glance, since human and gorilla show overexpression of both factors despite having smaller testes when compared to chimpanzee and gorilla. This becomes less surprising when taking the reported intricate relationships with their target genes into account: both MAX and TEAD1 appear to inhibit their targets’ expression when they are overexpressed alone, that is without their activating factors (MYC and YAP1/TAZ, respectively) in the above-mentioned contexts (Watt et al., 2017; Gu et al., 1993). It is therefore difficult at this point to deconvolute a putative mechanism that could have caused convergent testis size evolution observed between human-gorilla and chimpanzee-macaque. Nevertheless, our observations indicate the Hippo pathway as a candidate cellular mechanism behind convergent testis size and composition changes among primates.

We further studied putative regulators of Cluster #1 that do not show convergent expression evolution themselves, with respect to their roles in regulation of spermatogenesis and steroid hormone synthesis in response to the luteinizing hormone (LH). RFX1 and RFX2 were in this group: RFX2 has been reported to be the major regulator of spermatogenesis and forms a heterodimer with RFX1 (Wu et al., 2016; Kistler et al., 2015). Another factor, DLX5 regulates testicular steroidogenesis through GATA-4 (not enriched), which in turn binds to the *STAR* gene promoter to induce steroid synthesis (Nishida et al., 2008).

## CONCLUSION

Convergent evolution of testis anatomy has been previously linked to mating strategy differences among mammalian species. Here we show that the same convergent evolution patterns are traceable in bulk testis transcriptomes. We further report three main findings. First, these convergent transcriptome patterns are in large part explained by convergent tissue cell type composition shifts among species [as observed earlier among mice species (Good et al. 2010)]. Second, convergent changes at the cell-autonomous level can also be detected, especially among spermatids, but at modest levels compared to tissue composition changes. Thus, changes in ratios of already specialized cell types (PRE and POST) in response to relaxation or elevation of sexual selection appears as the main driver of testis transcriptome divergence patterns observed earlier among humans, chimpanzees and gorillas (Khaitovich et al., 2005; Brawand et al., 2011). Third, we demonstrate that human and gorilla testis transcriptomes are in paedomorphic states compared to those of chimpanzee and macaque (or the latter are hypermorphic compared to the former), mirroring the species’ anatomical differences. Change in the rate or timing of development, or heterochrony, could thus be one of the mechanisms behind the observed evolutionary shifts in tissue composition and gene expression.

Meanwhile, our study is limited by three main issues. One drawback involves being limited to only two single-male taxa, human and gorilla, with the latter represented by one individual. Despite this handicap, the clear parallels between relative testis mass distributions and transcriptome divergence patterns across these species supports the veracity of the convergent expression patterns we find, at least across the Old World anthropoids studied. We also note that non-primate species with manifest multi-male behavior, *Mus musculus* and *Rattus norvegicus*, have testis transcriptomes resembling those of multi-male primates. Future work may extend the analyses here to include additional species and test the generality of our observations. Importantly, this will require careful documentation of sampled individuals (including breeding season and status) and detailed information on species mating behavior for clear interpretation of the results (Methods).

A second weakness involves the use of indirect comparisons involving different data sets when inferring tissue composition and cell-autonomous changes in the testes. It would naturally be desirable to study these effects directly by conducting single cell transcriptomics experiments with multiple single- and multi-male primate testis samples. However, obtaining high quality postmortem material from most primates (except for humans and macaques) is extremely difficult and many times impossible. We therefore believe that reanalysis of existing datasets, as we have done here, is a valuable approach that helps address long-standing questions, and also guides future research.

A final drawback involves the restriction of our gene set to 1:1 orthologs, and the consequent exclusion of lineage-specific genes. This may lead us to overestimate the fraction of genes with convergent evolution signatures, as we cannot identify convergent changes among lineage-specific genes within our framework. Whether lineage-specific genes may also contribute to mating strategy-related phenotypic changes remains to be studied.

Despite these limitations, our results suggest that, as inter-male sperm competition escalates in a lineage, the testicles respond by boosting germ stem cell proliferation or extending its duration. This leads to hypergrowth and a higher proportion of seminiferous tubules, in addition to cell-autonomous expression changes in germ cells. Conversely, if inter-male competition subsides in a lineage, either the cost of sperm production or anatomical constraints may select for paedomorphic testicles, or the phenotype may simply drift towards a reduced phenotypic state by accumulation of disruptive alleles. We propose that the testis could be unique among other mammalian tissues in the rate of heterochronic changes leading to hypergrowth and paedomorphism, possibly by tinkering with the same key developmental regulators that yet await discovery.

## MATERIALS AND METHODS

Detailed information regarding preprocessing the raw data coming from the published transcriptome studies, merging and creation of combined datasets used in the analyses, and further discussion about species choice and phylogenies is available in the supplementary information.

### Differential expression and group differences

We used the non-parametric Mann-Whitney U (MWU) test (Wilcoxon rank-sum test) when testing for group differences in central tendency, applying the function “wilcox.test” from base R package “stats”. The MWU was chosen so as to be robust to outliers and skewed distributions. A two-sided MWU test was used to test each gene for differential expression (DE) between humans and chimpanzees, as well as in “Inter-species correlation analysis” and in “Developmental comparisons” (see below).

### Multiple testing correction

We corrected for multiple testing by converting p-values to q-values, using the Benjamini-Hochberg (BH) correction (Benjamini and Hochberg, 1995) as implemented in the function “p.adjust” from base R package “stats”.

### Inter-species correlation analysis

Here our goal was to determine whether an outgroup species’ transcriptome profile showed higher affinity to that of humans or to that of chimpanzees, treating these two as representatives for single- and multi-male mating, respectively. Under the null model of no mating strategy effect on testis transcriptomes, we would expect the expression profile of the outgroup species (i.e. gorilla, macaque, mouse, rat, marmoset, opossum, platypus) to be equally similar to those of human and chimpanzee. To increase resolution in the test, we limited the analysis to genes identified as differentially expressed (DE) between humans and chimpanzees. Across these DE genes, we calculated the Spearman correlation coefficient between (a) each outgroup species’ expression profile, (b) each human or chimpanzee individuals’ expression profile. If we had >1 individual for an outgroup species, for simplicity, we used the mean expression level across all individuals per gene to represent the species’ expression profile. For each species, we then tested for significant differences between its correlation coefficients to humans and to chimpanzees. For this, we used two approaches. First, we calculated p-values directly based on the MWU test. In the second, we used a permutation test approach, again using the Mann-Whitney U statistic. To obtain a p-value, we created a null distribution (N=10^5^) for each comparison by permuting the “chimpanzee” or “human” labels and recalculating the U values. Because we were to perform two-sided tests across different U value distributions, we centered the resulting null distribution for each comparison to zero by subtracting the mean of the distribution from the distribution itself, and shifted the observed U value using the same mean, for the sake of convenience during the p-value calculation described in the following. We finally calculated a two-sided p-value by comparing the absolute value of the observed U against the absolute value of the null distribution. The MWU test and permutation test p-values were essentially the same, and the latter is reported.

### Testing convergent evolution using the “EVE” model

We tested for convergent expression level evolution among primates with multi- and single-male mating systems using the combined primate dataset. Here we have a relatively closely related four-species phylogeny where neither the multi-(chimpanzee and macaque) and single-male (human and gorilla) taxa are monophyletic. Achieving the same phenotype related to mating strategy thus requires multiple mutational events, creating a powerful system to detect convergent evolution. To test for convergent expression level evolution for multi- and single-male species, we employed the Expression Variance and Evolution (EVE) method (Rohlfs and Nielsen, 2015), implemented as an R package (Gillard et al., 2020). EVE’s model-based approach accounts for phylogenetic relatedness among species while testing our alternative hypothesis of mating-type related convergent evolution. We compared the likelihood of the observed expression levels, (a) under a null model of stabilizing selection on expression levels pulling them towards a single optimum (M_0_ : θ_multi−male_ = θ_single−male_), versus (b) under an alternative model with one optimum for multi-male and another optimum for single-male mating systems (M_A_ ∶ θ_multi−male_ ≠ θ_single−male_). Since these models differ by one parameter, their likelihood ratio test statistics should be chi-square distributed with 1 degree of freedom under the null hypothesis of no convergent evolution. We thus calculated a p-value for each gene by comparing its likelihood ratio test statistic to the chi-square distribution using the R “stats” package function “pchisq”, and converted these further into q-values using the BH correction as described above. We used EVE to test the following datasets:

- Bulk testis transcriptome data for convergent evolution under mating strategy effects across 4 primates, i.e. between human-gorilla vs. chimpanzee-macaque;
- Bulk testis transcriptome data for convergent evolution between human-macaque vs. gorilla-chimpanzee (negative control for mating strategy across 4 primates);
- Bulk testis transcriptome data for convergent evolution under mating strategy effects across 3 mammals, i.e. between macaque-mouse vs. human;
- Bulk testis transcriptome data for convergent evolution between human-mouse vs. macaque (negative control for mating strategy across 3 mammals);
- Spermatocyte or spermatid transcriptome data for convergent evolution under mating strategy effects between mouse-macaque vs. human;
- Spermatocyte or spermatid transcriptome data for convergent evolution between mouse-human vs. macaque (negative control for mating strategy);
- Brain transcriptome data for convergent evolution under mating strategy effects across 4 primates, i.e. between human-gorilla vs. chimpanzee-macaque;
- Brain transcriptome data for convergent evolution between human-macaque vs. gorilla-chimpanzee (negative control for mating strategy);
- Predicted cell type proportion (log_2_(b_POST_/b_PRE_) ratios, see below) using primate data;
- Predicted cell type proportion (log_2_(b_POST_/b_PRE_) ratios, see below) using primate and rodent data.

### Cell type proportion prediction

Here we predicted the relative contribution of pre-meiotic/somatic (PRE) versus meiotic/post-meiotic (POST) cell types to the overall adult testis transcriptomes using a deconvolution-like approach (Gong et al., 2011). First, for each of the n=7,135 genes in the combined mouse cell type dataset, we calculated mean expression levels for all PRE (n=8) and POST (n=8) transcriptome samples, denoted by E_PRE_ and E_POST_, respectively. Of these genes, n=4,724 were common with the adult mammal bulk testis data. Next, using gene expression values across bulk testis samples, denoted E_WT_, we constructed a linear regression model across all overlapping genes:

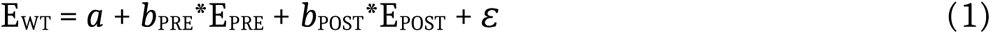

where *a* represents the intercept, *b*_PRE_ and *b*_POST_ represent the regression coefficients, and ε stands for error. Using these coefficients, we then calculated the ratio log stands for error. Using these coefficients, we then calculated the ratio log_2_(*b*_POST_ / *b*_PRE_) for each individual, which estimates the relative contribution of POST vs. PRE cells to the bulk tissue sample (Figure 2A).

### Role of tissue cell type composition shifts in convergent transcriptome changes

Here our goal was to test whether tissue composition shifts explain convergent expression divergence. For this, we used the information from the combined primate dataset (adult bulk testis transcriptomes) and the combined mouse cell type dataset. There were n=6,738 genes common between the two datasets. For all common genes, we calculated two effect size measures, using the Cohen’s D statistic with pooled variance (Lakens, 2013):

- PRE versus POST expression effect size in the mouse cell type dataset, which we denote E_PRE-POST_;
- Single-male versus multi-male expression effect size in the combined primate dataset, which we denote E_SM-MM_.

These were calculated separately for each gene. We then calculated the Spearman correlation between E_PRE-POST_ and E_SM-MM_ across the common genes (ρ = 0.40 p<10 = 0.40 p<10^-15^). We further compared the absolute E_PRE-POST_ values between genes showing evidence for convergent evolution in the EVE analysis (at q<0.10), and genes showing no such indication (at q>0.90). Compared to genes showing no convergent evolution, convergently evolving genes had 1.33 times higher median absolute E_PRE-POST_ (MWU p<10^−7^).

### Bulk testis vs. cell-autonomous divergence

Here our goal was to study how much whole (bulk) testis transcriptome divergence could be explained by cell-autonomous (cell-specific) expression changes. We used two approaches, described in the following.

### Bulk testis vs. cell-autonomous divergence studied via multiple regression

We aimed to estimate the contributions of cell type shifts and cell-autonomous changes to bulk testis divergence between single- and multi-male species, using multiple regression models. We studied 3 multiple regression models; they had the same explanatory factors, but different responses. Both responses and explanatory variables were effect size values [Cohen’s D with pooled variance (Lakens, 2013)] calculated separately for each of the n=4651 common genes among datasets.

- Responses: these were effect size values calculated between single- and multi-male species pairs from bulk testis (BT) data. The pairs were human-macaque, human-chimpanzee, and gorilla-chimpanzee, denoted E_hsa-mml(BT)_, E_hsa-ptr(BT)_, E_ggo-ptr(BT)_, respectively.
- Explanatory factor 1: this was pachytene spermatocyte (PS) expression effect size between human-macaque, denoted E_hsa-mml(PS)_.
- Explanatory factor 2: this was round spermatid (RS) expression effect size between human-macaque, denoted E_hsa-mml(RS)_.
- Explanatory factor 3: this was PRE versus POST expression effect size in the mouse cell type dataset, denoted E_PRE-POST_.

The models were the following:

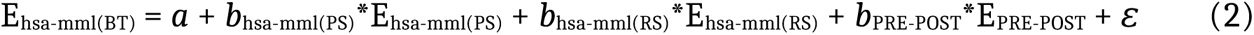

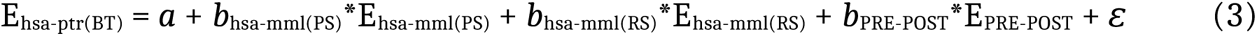

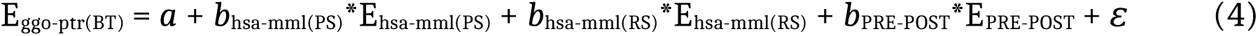

where *a* represents the intercept, *b*_hsa-mml(PS)_, *b*_hsa-mml(RS)_, *b*_PRE-POST_ represent the regression coefficients, and ε stands for error. Using these coefficients, we then calculated the ratio log stands for error.

We expect that:

- If cell-specific transcriptome divergence contributes to bulk testis divergence, *b*_hsa-mml(PS)_ and/or *b*_hsa-mml(RS)_ may be valid estimators of E_hsa-mml(BT)_, and possibly of E_hsa-ptr(BT)_.
- If cell-specific transcriptome divergence contributes to *convergent evolution* of bulk testis expression patterns, then *b*_hsa-mml(PS)_ and/or *b*_hsa-mml(RS)_ may be valid estimators of all responses, including E_ggo-ptr(BT)_.

### Bulk testis vs. cell-autonomous divergence studied via branch-length correlations

For this, we used single male species’ expression divergence in species trios that included the following taxa: (a) a single-male primate, (b) a multi-male primate, (c) a multi-male rodent species.

For the bulk testis data, we used 3 trios:

- Human-rhesus-mouse,
- Human-chimpanzee-mouse,
- Gorilla-chimpanzee-rat.

For the cell-specific data there was only one trio possible: human-rhesus-mouse (Figure 2B).

For each trio, for each gene, we constructed a neighbor-joining tree using the R “ape” package function “nj” (Paradis and Schliep, 2019). For each such tree, we then calculated the single-male species’ branch length, which we abbreviate as SMBL.

We tested correlation between bulk testis SMBL and cell-specific SMBL values (across n=4894 common genes) using Spearman correlation.

We expect that:

- If cell-specific transcriptome divergence contributes to bulk testis divergence, cell-specific SMBL and bulk testis SMBL should be correlated, at least for SMBL that include common species (e.g. cell-specific SMBL from human-rhesus-mouse and bulk testis SMBL from human-chimpanzee-mouse).
- If cell-specific transcriptome divergence contributes to *convergent evolution* of bulk testis expression patterns, then cell-specific SMBL and bulk testis SMBL should be correlated across all trio pairs, as long as the mating strategy-related topology remains fixed (e.g. cell-specific SMBL from human-rhesus-mouse and bulk testis SMBL from gorilla-chimpanzee-rat).

The analysis was performed separately for cell-specific spermatocyte and spermatid datasets.

### Developmental comparisons

Our goal here was to understand whether closely related single- and multi-male primate species, and in particular human and chimpanzee, differ in their developmental trajectories of the testis using transcriptome data. We further aimed to test the hypothesis that single-male adult testis transcriptome profiles could be paedomorphic relative to those of multi-males species, in parallel with their anatomy. Because chimpanzee testis development series are unavailable, we used an indirect approach, and compared adult bulk testis transcriptome profiles to transcriptome profiles from mouse and macaque developmental series. We chose these two multi-male species because, under the hypothesis of single-male paedomorphosis, an adult multi-male species’ expression state should surpass that of an adult single-male species (e.g. for a gene that increases in expression level during development, adult chimpanzee expression should be higher than that of adult human).

We first used the mouse testis development dataset described above, with n=15 mice with ages ranging from postnatal day 0 to 60. We applied following procedure:

- We identified n=2,295 genes showing significant expression level change with mouse age using the Spearman correlation test (at q<0.10).
- For each of these 2,295 genes, we interpolated loess regression curves of expression change with age using the “loess” function in the R “stats” package, across 60 points (from age 1 to 60 days), and with the degree parameter fixed at 1 (note that interpolation allows studying expression changes uniformly across development, without being limited to the ages of the sampled mice).
- We calculated Spearman correlation coefficients between these interpolated mouse testis expression levels, and each adult primate species’ testis profile, as well as the PRE and POST profiles, at each interpolated point, across all 2,295 common genes. We performed this analysis using the mean expression level for each species and each cell type.

The correlation coefficients were plotted against mouse age. To better observe differences among correlation coefficient curves, we further normalized each set of correlation coefficients to mean=0, and s.d.=1.

To test for human paedomorphism, i.e. whether the human curve is significantly shifted to earlier ages compared to the chimpanzee curve, we applied following procedure:

- We again calculated correlation coefficients between the interpolated mouse profile and each hominid species.
- For both species, we determined the mouse age at which the correlation is maximized, or the “age of maximum expression similarity”.
- We tested whether the age of maximum expression similarity is lower for humans than for chimpanzees, using a one-sided permutation test approach, and using the Mann-Whitney U statistic, similar to the permutation test applied in the correlation analyses described above.

The same steps were repeated to analyse the macaque testis development dataset, which includes n=12 rhesus macaques with ages ranging from postnatal day 16 to 26 years. We used log2-transformed age (in days) to more efficiently model expression changes during development, which are fast early in life but slower later (>4 years). This dataset contained n=2,321 development-related genes.

The same steps were repeated to analyse the mouse neocortex development dataset, which includes n=8 mice from postnatal day 2 up to 122 days of age. This dataset contained n=3,314 development-related genes. We used the human, chimpanzee, gorilla and rhesus macaque prefrontal cortex profiles to compare with mouse neocortex development.

### Gene clustering

Here our goal was to identify genes with common patterns throughout human, macaque and mouse testis development, and further to study their functional enrichment, their enrichment among convergently evolving genes, and their regulation. We clustered genes solely based on testis development profiles across these 3 species using the k-means algorithm. Specifically, we applied the following procedure:

- In all 3 testis development datasets, we scaled each genes’ expression to mean=0, and s.d.=1. Scaling removes the influence of average expression level differences among genes, and maximises information on variation among samples.
- We merged the 3 scaled datasets. The resulting dataset contained n=5,596 common genes.
- We clustered the merged dataset using k-means. For this we first searched for the optimal number clusters (k) using the gap statistic. This is implemented in the “clusGap” function in the R base package “cluster”. This did not yield a practical result in that it either yielded an unreasonably high k (e.g. 26), or a reasonable k (e.g. 13) but then k-means runs did not converge. For this reason, we also studied the information captured in average silhouette scores for k={4,..,14}. k=6 appeared as a good trade-off between having a too small k (having too few clusters with large membership could lead to merging of unrelated pathways and make downstream functional analyses harder) and a too large k (too many clusters with too few members could lead to inconclusive downstream functional analyses).
- Because k-means is a heuristic algorithm we increased the maximum iterations to 500 and started with 500 random initial centroids to obtain the most optimal clustering.

We also confirmed that (a) the overall expression variation patterns we see with 6 clusters, and (b) the spermatogenesis-related GO term enrichment in the cluster with overall increasing testis development expression and adult multi-male overexpression among convergent genes were consistently found for each k tested (Figure S11).

### Enrichment for convergent genes

We tested genes in each of the 6 clusters for enrichment for genes showing convergent expression patterns with respect to mating strategy (at q<0.10, according to the EVE results), relative to genes not showing convergent expression patterns (at q>0.10). We used genes in all other clusters as background, and implemented the Fisher’s exact test (“fisher.test” in the R “stats” package). Clusters #1 and #6 showed such enrichment.

### Gene Ontology enrichment

We tested convergent genes in Clusters #1 and #6 for enrichment in Gene Ontology (GO) (Ashburner et al., 2000) Biological Process categories. For this we used the “TopGO” R package (Alexa and Rahnenfuhrer, 2019), with the “parentChild’’ algorithm for the evaluation of contingency tables, which is suggested to reduce false-positives resulting from the hierarchical structure of the GO (Grossmann et al., 2007). We used BH correction on the Fisher’s exact test p-values calculated by “TopGO”. When testing each cluster for GO enrichment, convergent genes in that cluster used as the foreground and the non-convergent genes that are not from that particular cluster were used as the background. This makes the tests biologically relevant to our hypotheses, testing convergent genes that share a common developmental pattern against genes that are not in this developmental “module” *and* are not convergent.

### Transcription factor binding site enrichment

Here our goal was to study the regulation of genes in Cluster #1. We used TRAP and PASTAA (Roider et al., 2007, 2009) for calculating binding affinities of transcription factors (TF) and enrichment in Cluster #1 genes, respectively. Positional Weight Matrices for transcription factors were retrieved from the “MotifDb” R package (Shannon and Richards, 2019). Human sequences 2 kb upstream from each gene’s transcription start site were retrieved using the “biomaRt” R package (Durinck et al., 2009). PASTAA and TRAP source code was downloaded from http://trap.molgen.mpg.de/PASTAA/. When run with a discrete set of foreground and background genes, PASTAA optimizes the number of targets among the foreground and among all genes by iteratively performing hypergeometric tests (HT) with changing numbers of genes labeled as targets for both gene sets using the TF and gene affinity information coming from the output of TRAP. Number of genes labeled as targets of a given TF among the two categories that yields the most significant HT p-value reported as the optimal number of targets. The background gene set was chosen as the intersection of all non-convergent and all genes outside cluster #1, using the same approach as described above for GO enrichment. MotifDb is a comprehensive database supplied as an R package and it contains Positional Weight Matrices from several sources. To de-duplicate the results after enrichment, resulting enrichment scores were filtered such that we retained only one Positional Weight Matrix per TF. This was achieved by selecting for the entry with the lowest raw enrichment p-value among several matrices for a given gene. De-duplicated entries were then corrected for multiple testing using the BH approach as above. Top targets of selected TFs represented in Figure S10 obtained for each TF by using the reported optimal number of targets (N) among all genes used and selecting top N genes to which the given TF show the most affinity to according to the TRAP output.

## Author contributions

(a) M.S., P.K. and R.R. conceived and designed the study; (b) Z.Y. performed the laboratory experiments; (c) E.Y., E.S., M.D., performed the data analyses with contributions from H.H., S.G., E.O., M.S., R.R., B.E.; (d) M.S. and R.R. supervised the analyses; (e) M.S. and E.Y. wrote the manuscript with contributions from E.S., R.R., M.D., and E.O., and all authors approved the final version of the manuscript.

## Data accessibility

All raw data created is accessible on NCBI GEO database. Accession numbers GSE73636 and GSE73635 can be used to access macaque testis development and mouse brain development datasets, respectively.

## Acknowledgements

We are grateful for Iris Finci for sharing data, Duha Alioglu, Ayşegül Dede and all members of the CompEvo Group at METU, members of the Comparative Biology group at Shanghai, Tülin Cetin, Mesut Muyan, Henrik Kaessman, Marie Semon, Ömer Gökçümen, Rafik Neme, Jamie Winternitz, and two anonymous reviewers for discussion and/or suggestions on the manuscript. We thank Tülin Yanık for support. M.S. was supported by a TUBITAK 2232 fellowship (no. 114C040) and a Science Academy-Turkey Award (BAGEP-2014), and METU BAP-07-02-2015-009. The funders had no role in study design, data collection and analysis, decision to publish, or preparation of the manuscript. All species icon illustrations were downloaded as royalty free vector images from various websites such as freepik.com, vectorstock.com, and vecteezy.com.

## Supplementary Figures

**Figure S1:**
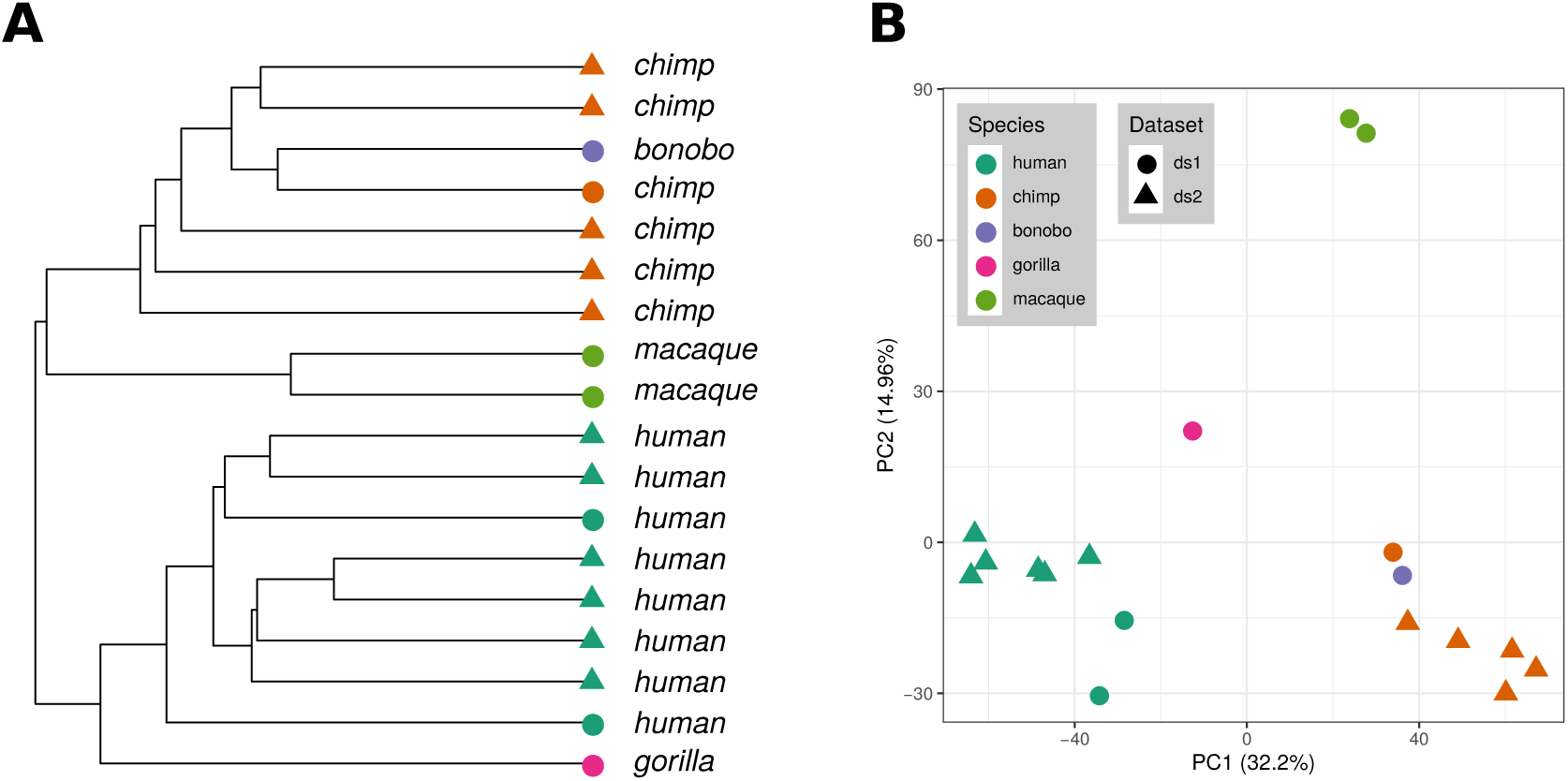
Quality control and overall inspection of the combined primate testis dataset.**(A)** UPGMA hierarchical clustering dendrogram of all individuals. **(B)** Principal components (PC) analysis plot of the first two PCs. The percent of variance explained by each PC is shown on the axis labels. Labels ds1 and ds2 correspond to primate datasets no. 1 (**Brawand2011**) and no. 2 (**Khaitovich2005**), respectively.

**Figure S2:**
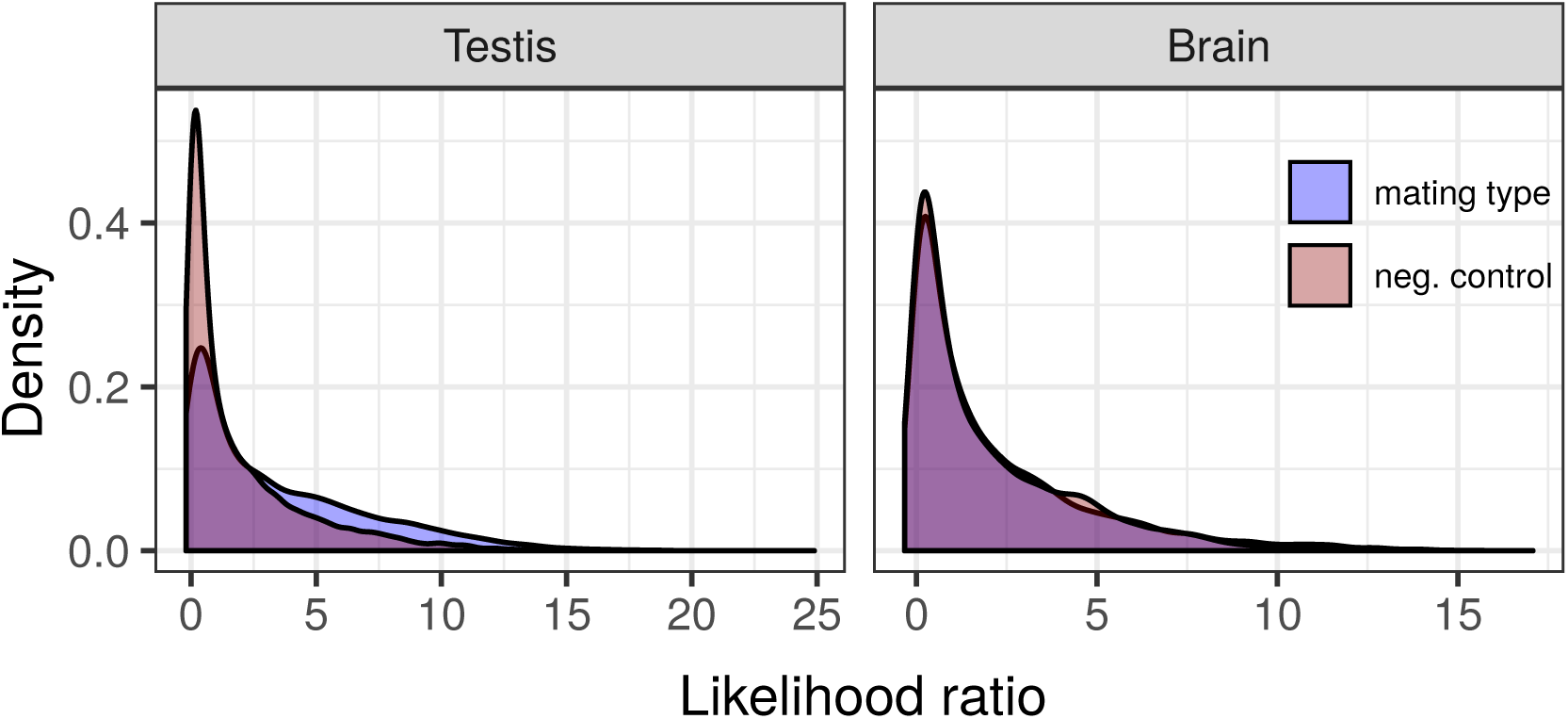
Likelihood ratio (LR) distributions of testis and brain EVE runs. Blue shaded areas show the LR distribution for the mating strategy-related convergent evolution tests whereas the red shaded areas show those of negative control tests, performed using the 4 Old World anthropoid species, as in Figure 1C.

**Figure S3:**
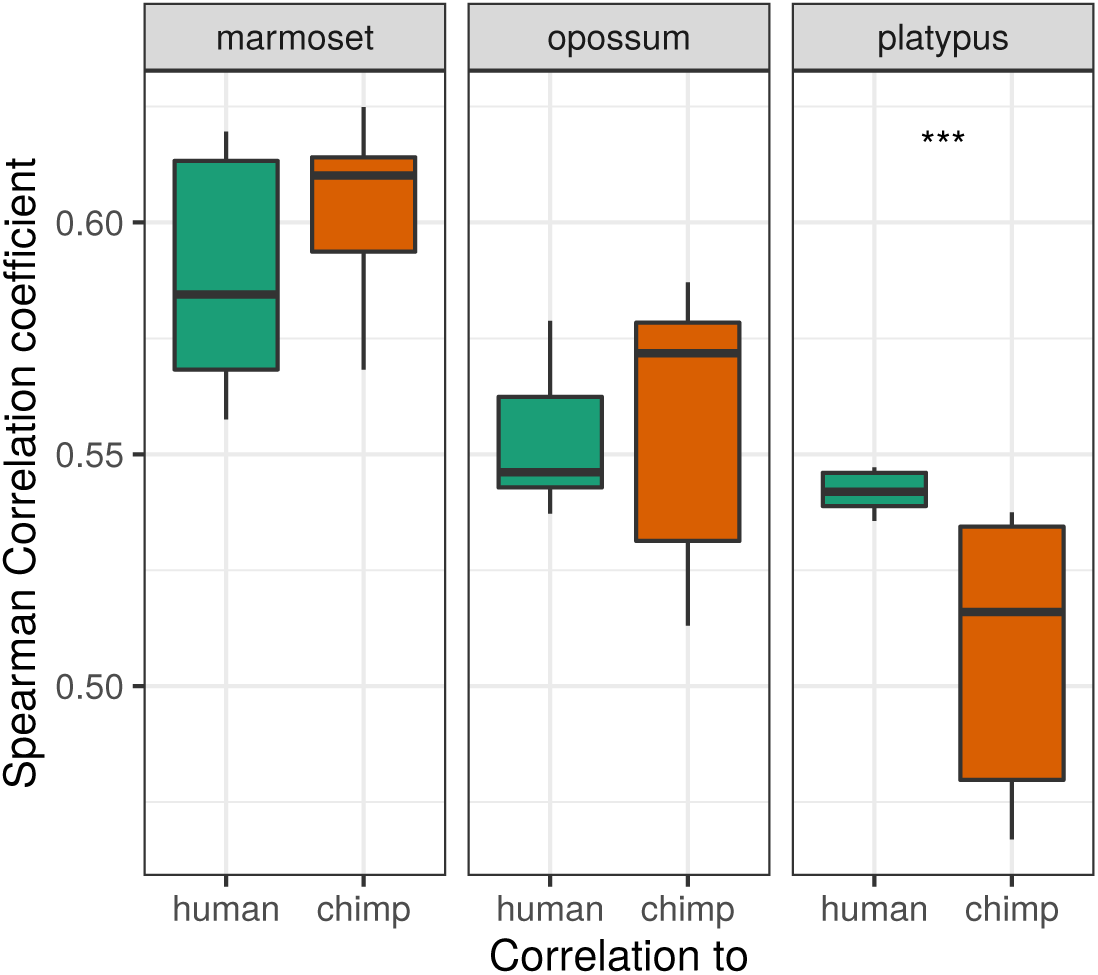
Correlation boxplots for additional mammalian species, calculated as in Figure 1B. The platypus is the only one among the three to show a significant difference in its relative correlation to humans vs. chimpanzees (permutation test, *p <* 10*^−^*^3^).

**Figure S4:**
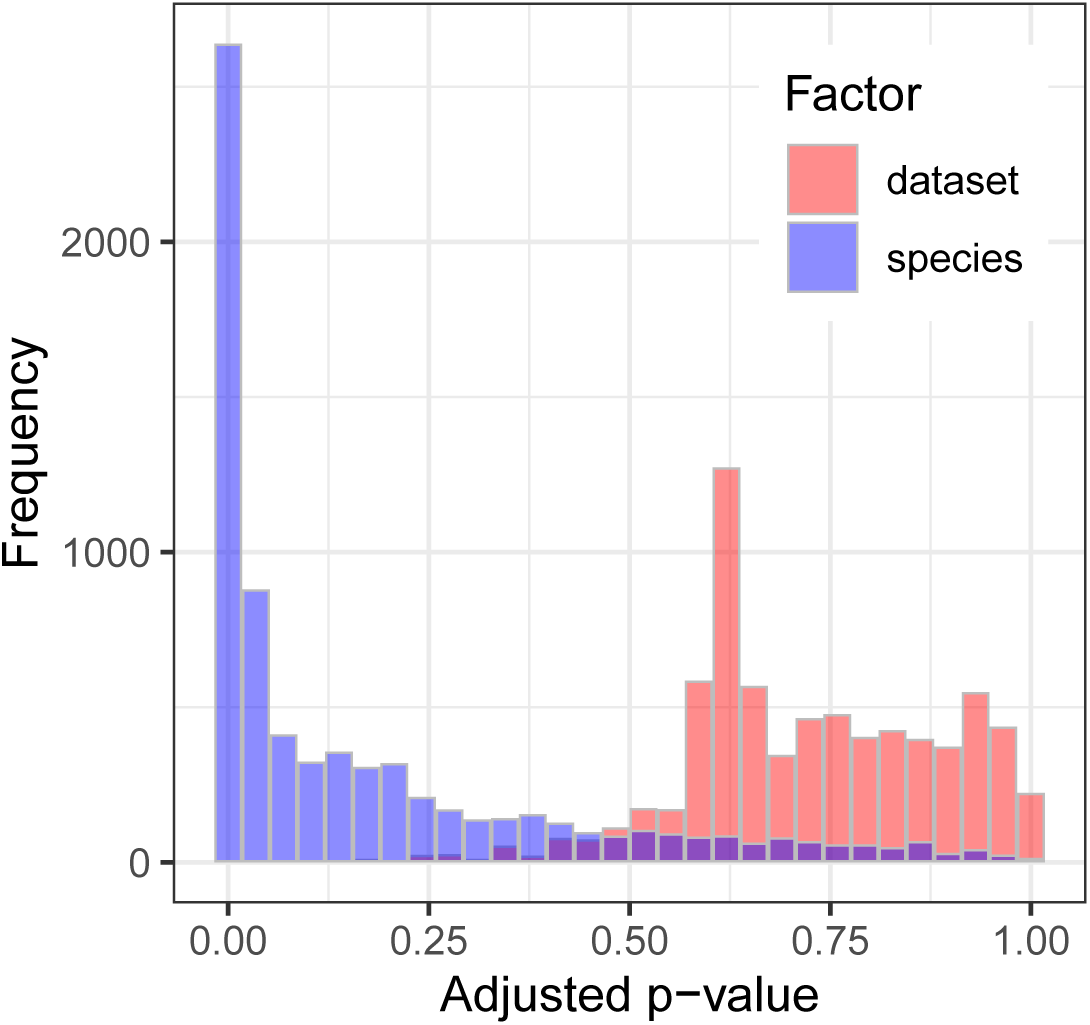
Adjusted p-value distributions of two-way ANOVA runs across n=7,305 genes present in the combined adult primate bulk testis dataset showing the effect of dataset source and species for each gene.

**Figure S5:**
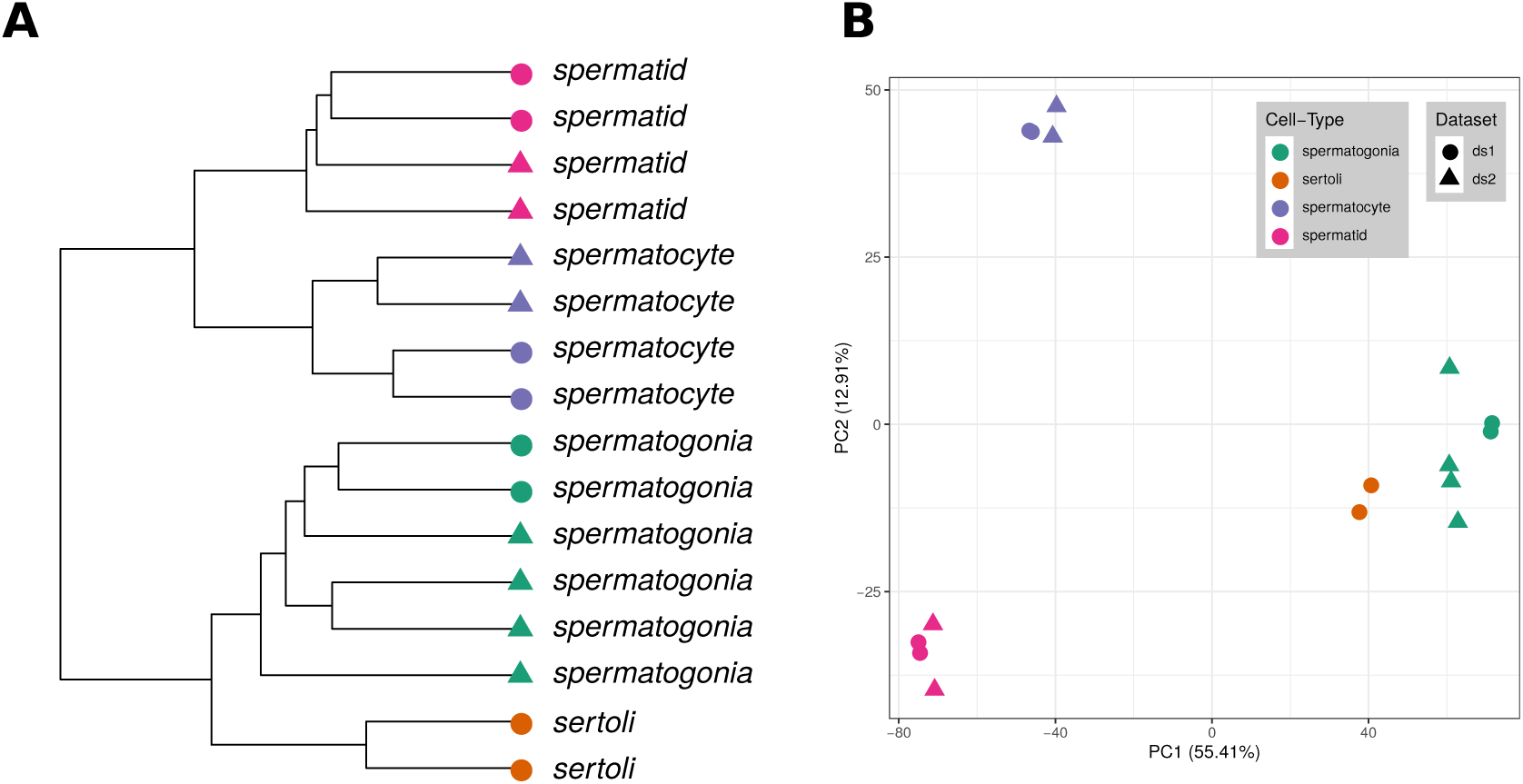
Quality control and overall inspection of the combined mouse cell-type dataset. **(A)** UPGMA hiearchical clustering of all 8 cell type samples. **(B)**Principal Component Analysis, plot of first two PCs. The percent of variance explained by each PC is shown on the axis labels. Labels ds1 and ds2 correspond to mouse cell-type datasets (**Chalmel2007**) and (**Namekawa2006**).

**Figure S6:**
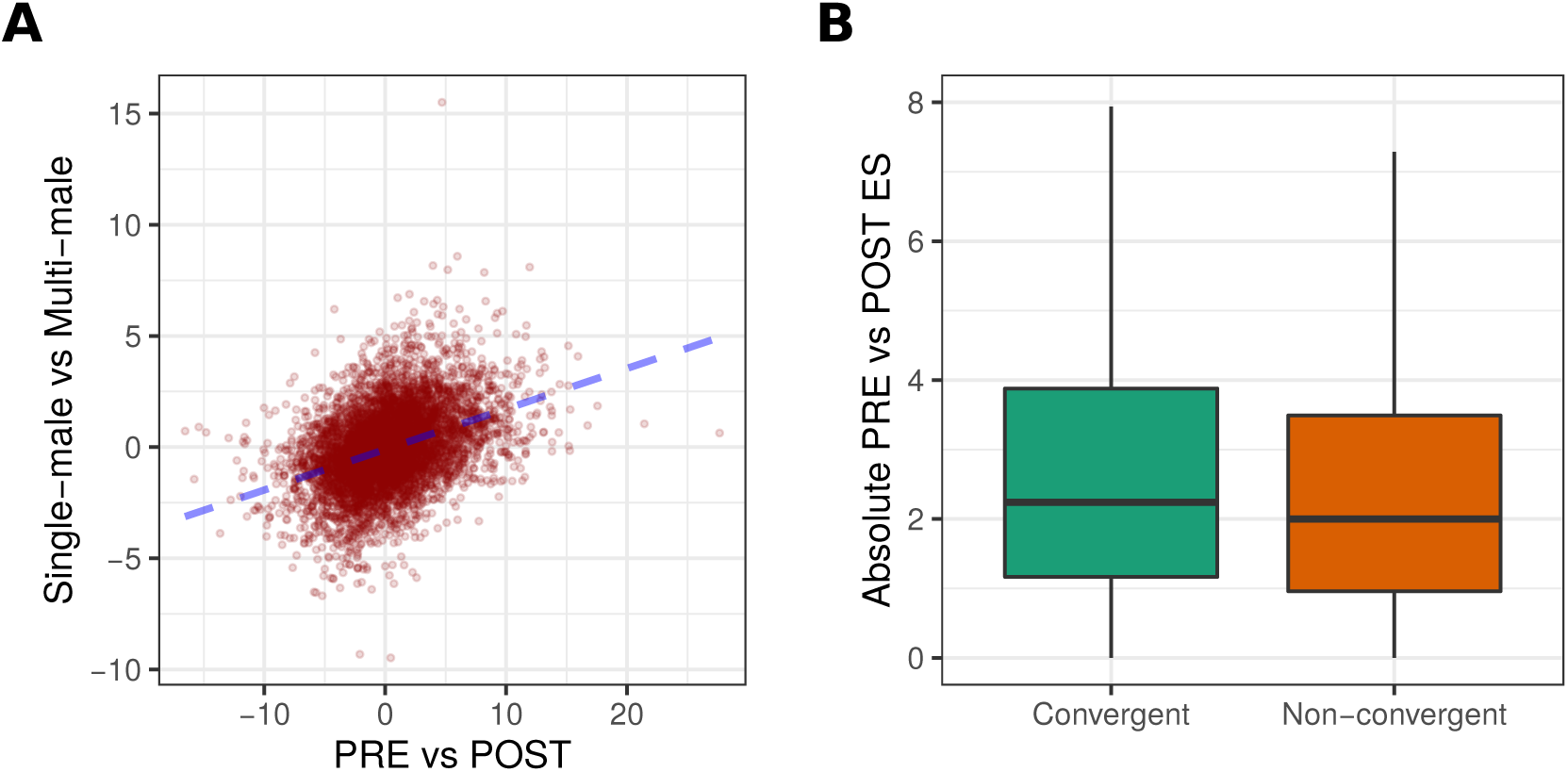
Measuring the contribution of tissue cell type composition changes to convergent evolution. **(A)** The y-axis shows effect sizes calculated between single-male (human-gorilla) versus multi-male (chimpanzee-macaque) species bulk testis transcriptomes; the x-axis shows effect sizes calculated between PRE versus POST cell type profiles. The correlation was calculated across n=6738 genes; Spearman correlation rho = 0.40, *p <* 10*^−^*^15^. **(B)** Distribution of absolute PRE versus POST effect size calculated for genes showing evidence of mating strategy-related convergent evolution in the EVE test (between human-gorilla versus chimpanzee-macaque; at *q <* 0.10; n=1970 genes) and calculated for all other genes (at *q >* 0.90; n=4768 genes, also see Figure 1C-D). Convergent genes have significantly higher absolute cell type effect size (one-sided Mann-Whitney U test, *p <* 10*^−^*^7^).

**Figure S7:**
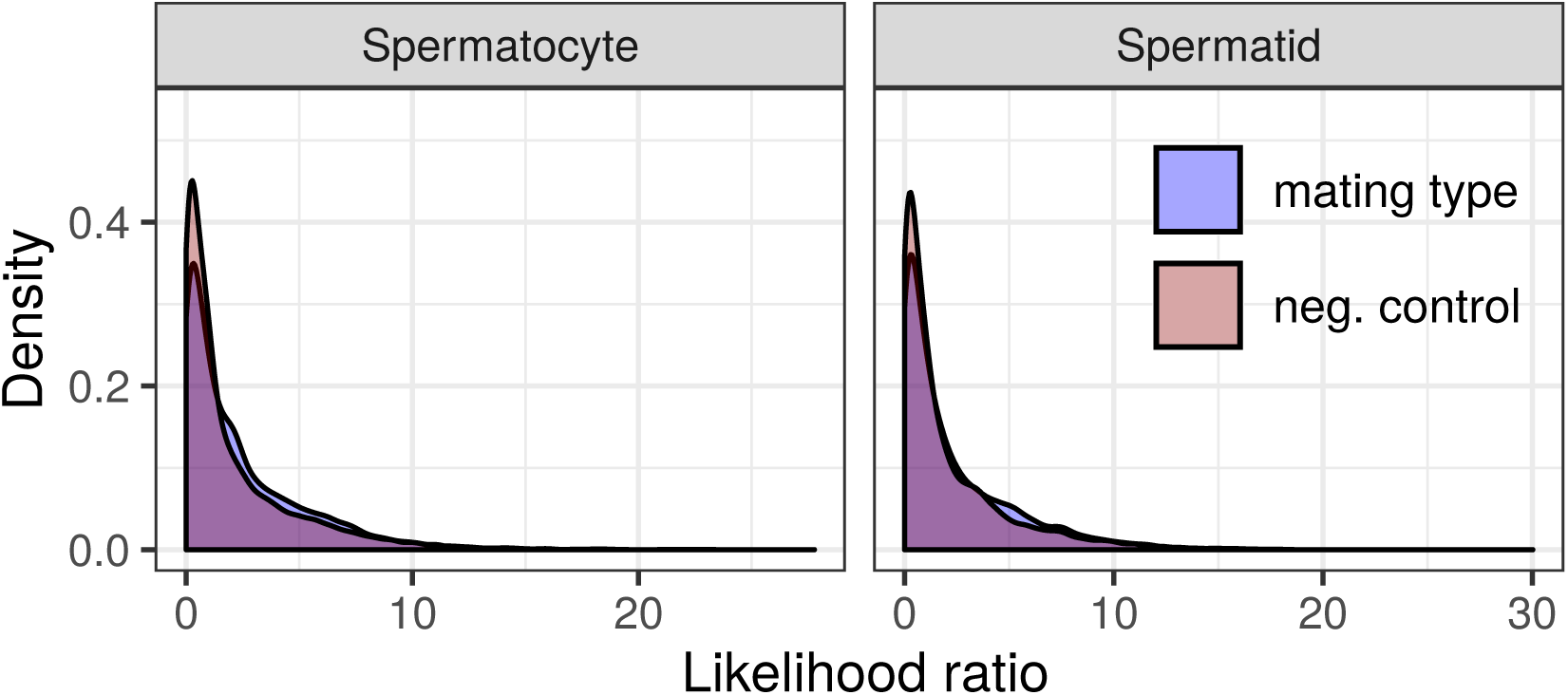
Likelihood ratio (LR) distributions of spermatocyte and spermatid EVE runs. Blue shaded areas show the LR distribution for the mating strategy-related convergent evolution tests [(macaque-mouse) vs (human)] whereas the red shaded areas show those of negative control tests [(human-mouse) vs (macaque)].

**Figure S8:**
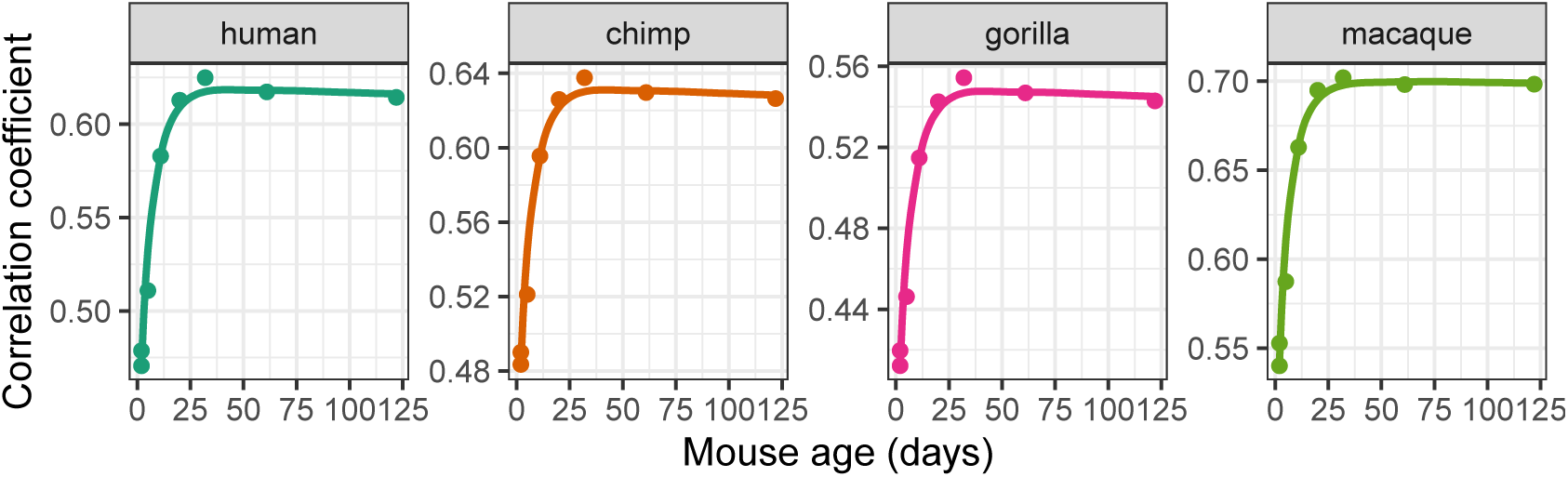
Correlations between adult primate brain (prefrontal cortex) expression profiles and mouse neocortex developmental profiles. The analysis was performed in the same way as in Figure 3. Across mouse neocortex development, both the human and gorilla (median=57 days of age) and the chimpanzee and macaque (median=43 days) show the highest correlation to adult mice. Although the median time-point to which human and gorilla show the maximum correlation across mouse neocortex development is nominally later than those of chimpanzee and macaque, this difference is not statistically significant (permutation test using Mann-Whitney U statistic, *p* = 0.81)

**Figure S9:**
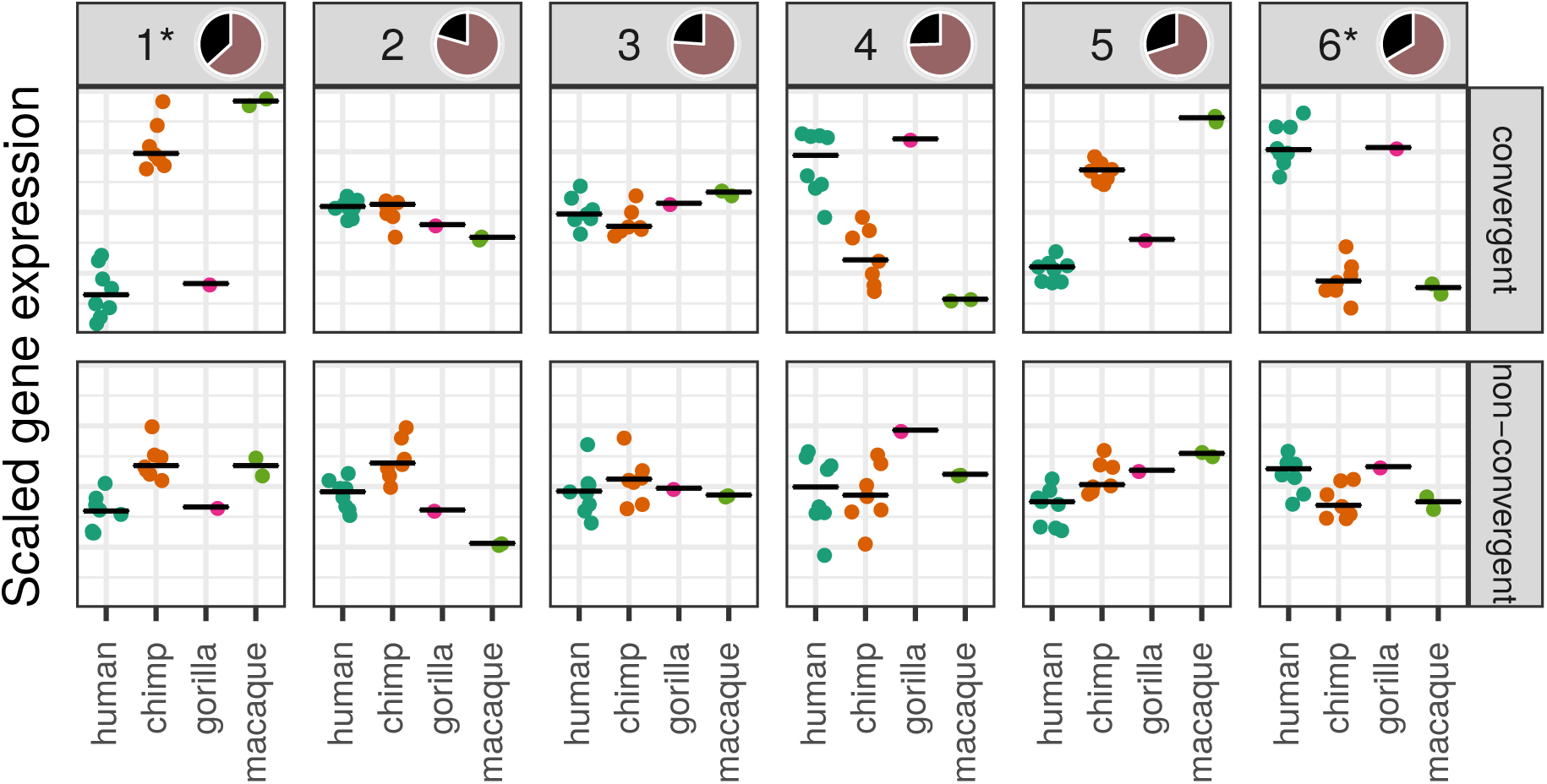
Expression variation among adult primates across developmental gene clusters. The y-axis shows mean gene expression levels of adult primate bulk testis samples across the same 6 gene clusters shown in Figure 4. The expression levels of genes are scaled to mean 0 and standard deviation 1 before calculating the means. For each cluster, the top row panes show mean expression of genes identified as convergently evolving in EVE analysis across the 4 Old World anthropoids, while the bottom row panes show non-convergent genes (i.e. all other genes) in that particular cluster. Relative abundances of convergent vs. non-convergent genes for each cluster shown as pie charts in pane labels. The dark portions correspond to the proportion of convergent genes, while the light portions correspond to that of non-convergent genes. **(*):** Clusters enriched in convergent genes.

**Figure S10:**
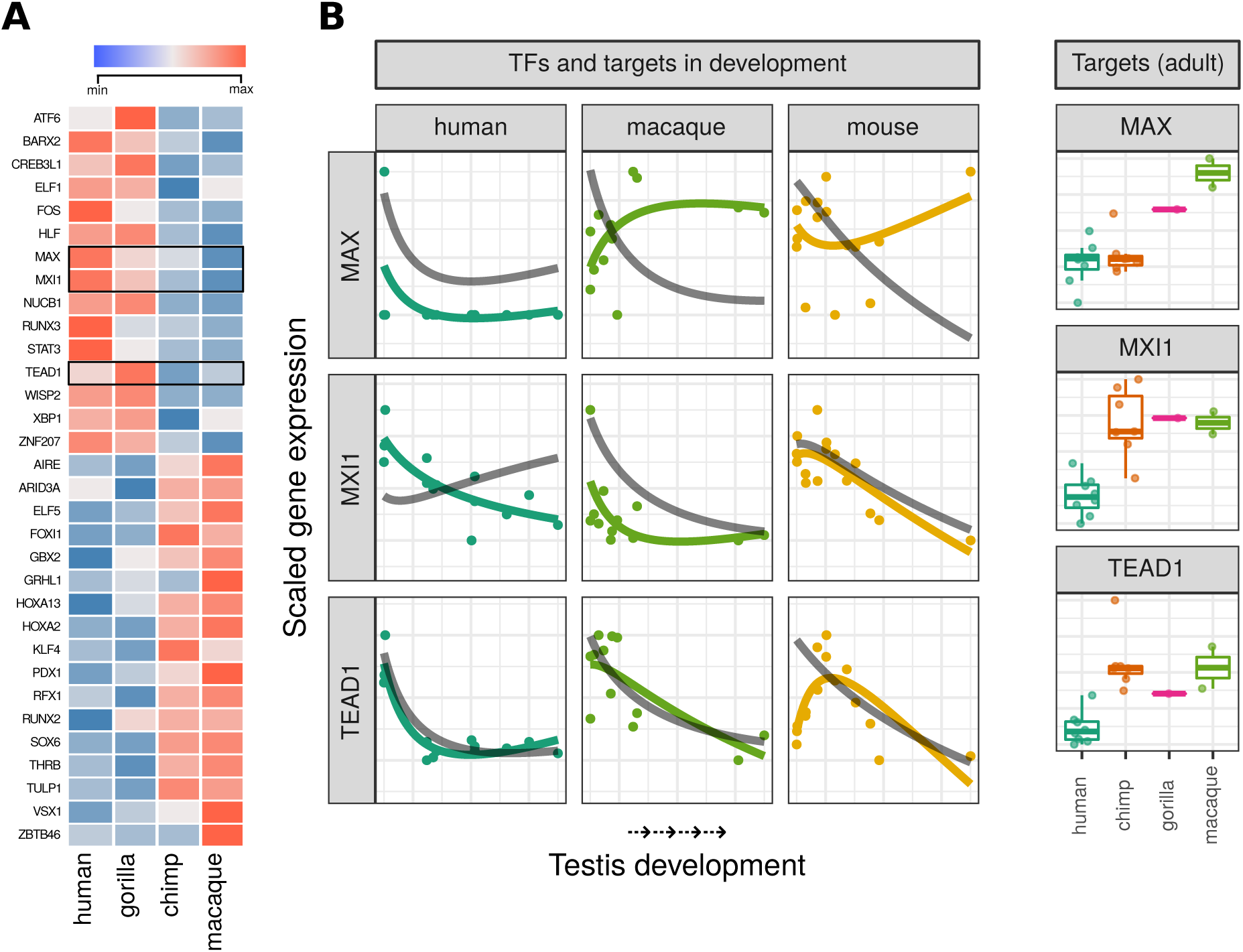
Putative transcription factor (TF) regulators of Cluster #1. **(A)** Heatmap of adult primate bulk testis gene expression levels of the 32 TF that are found enriched in Cluster #1 targets and that are also identified as convergently evolving in EVE analysis across the 4 Old World anthropoids. Each gene’s expression levels were scaled across the 4 species so that differences between species across genes are equally visible. **(B)** Left panels: Developmental trends of selected TF and their targets through testis development in human, macaque and mouse. The x-axis shows age, from newborns/infants to adults of each species, while the y-axis shows scaled gene expression. Colored lines represent gene expression levels of the given TF; gray lines represent the median gene expression of top targets of the given TF. For each curve, expression levels of genes are scaled to mean 0 and standard deviation 1 before plotting and/or calculating the medians. Curves are obtained using loess models constructed with the formula *y ∼ log*_2_(*x*) where y represents the gene expression level, and x represents the time-points throughout development. Right panels: Adult bulk testis median gene expression profiles of the same targets of the selected transcription factors. Median expression values were scaled before visualization. Genes labeled as the top targets of a given TF are selected by utilizing the TRAP and PASTAA output (See Methods for detailed explanation).

**Figure S11:**
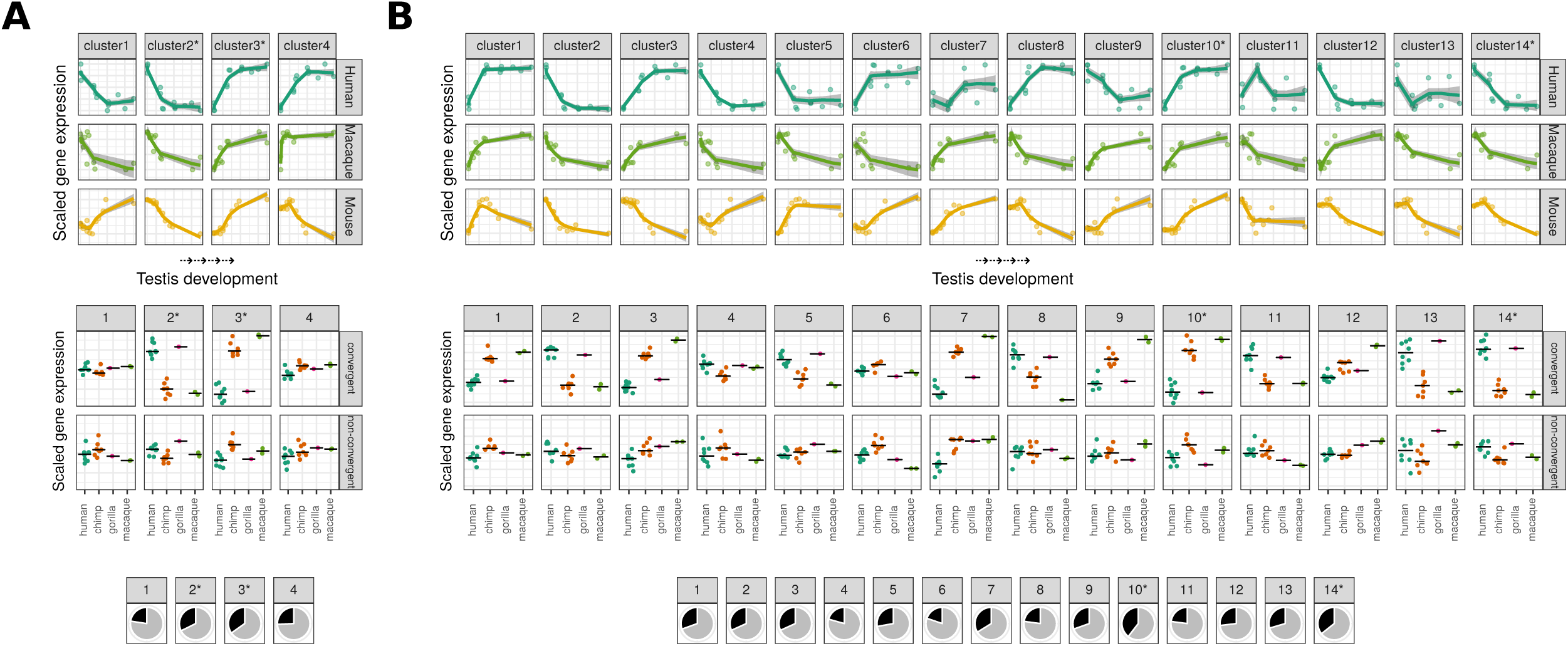
Reproduction of the Figure 4 and Figure **??** for clustering results obtained by setting **(A)** k=4 and **(B)** k=14.

