## Supplementary Information for "Convergent evolution of primate testis transcriptomes reflects mating strategy"

### Novel transcriptome datasets

**Ethics statement:** Samples used for microarray analysis in this study were derived from animals that died of reasons independent of this work. The Biomedical Research Ethics Committee of Shanghai Institutes for Biological Sciences completed the review of the use and care of the animals within the research project (approval ID: ER-SIBS-260802 P).

**Macaque testis development:** Post-mortem testis samples from n=12 rhesus macaques of different ages (16 days to 26 years) were obtained from the SuZhou Experimental Animal Center (SuZhou, China). The individuals had been housed under standard conditions, were healthy, and died of causes with no relation to the tissue used, and of reasons unrelated to this study. Whole testis samples were stored at -80°C. Total RNA from 100 mg tissue was isolated using the Trizol reagent. Biotinylated cDNA were prepared from 1 micrograms of total RNA, and hybridized to Affymetrix Human Gene 1.0ST arrays following standard Affymetrix protocols. The raw data is accessible through the NCBI GEO database (www.ncbi.nlm.nih.gov/geo/) under accession number GSE73636. We determined Affymetrix microarray probes that matched the rhesus macaque genome perfectly and uniquely, by aligning probe sequences to the rheMac2 genome sequence using BLAT (Kent, 2002). Affymetrix CEL files were processed using only those probes that fulfilled this criteria, following (Somel et al., 2010). Briefly, intensities of the chosen probes were compared with those of antigenomic probes of the same GC content, which represent background noise. Only those probes with intensity >95% of the background were included in further analysis. Probe intensities were then log-transformed, quantile normalized across arrays, and intensity values per probeset were calculated by median polishing, using the R "affy" package (Gautier et al., 2004). Per individual, a probeset was considered "detected" if >50% of probes and at least 8 probes passed the described filters, per probeset. We considered a probeset as "expressed" if it was detected in >70% of individuals. Affymetrix probeset IDs were mapped to human Ensembl genes following the procedure described in Section 2 of Materials and Methods.

**Mouse brain development:** Post-mortem neocortex samples were obtained from n=12 C57BL/6 mice aged between 2 days and 904 days, housed and fed under standard conditions in the MPI-EVA Animal Facility (Leipzig, Germany). The mice were sacrificed for reasons independent of this study. Cortical tissue was harvested and frozen immediately, and stored at -80°C. We extracted total RNA from supplied frozen prefrontal cortex tissue, using the Trizol reagent. RNA was processed and hybridized to Affymetrix Mouse Gene 1.0ST arrays following standard Affymetrix protocols. The raw data is deposited to NCBI GEO under accession number GSE73635. Affymetrix CEL files were processed using the Bioconductor “oligo” package “rma” function (Carvalho and Irizarry, 2010). Affymetrix probeset IDs were mapped to Mus musculus genes and further to orthologous human Ensembl genes following the procedure described in Section 2 of Materials and Methods. To create the mouse brain development dataset, we included data from n=8 mice from 2 days up to 122 days of age (excluding old adult mice).

### Published datasets and data preprocessing

***Custom transcriptome annotation (GTF) files including 1:1 orthologous exon sets***: Gene expression quantification using RNA-Seq data is alignment-based, and is therefore affected by the length of the annotated genes and exons. Since we deal with data obtained from several species, it was crucial to address possible differences in annotation and gene content completeness across different genomes. To achieve this we created two sets of custom transcriptome annotation (GTF) files that included 1:1 orthologous and perfectly alignable exons:

- GTF file A: This includes only the primate species analyzed in the study (human, chimpanzee, gorilla, macaque, marmoset),
- GTF file B: This includes all the mammalian species analyzed in the study (the aforementioned primate species, mouse, rat, opossum, platypus).

For this, we first collected sets of 1:1 orthologous genes from Ensembl v83 (Cunningham et al., 2015; Yates et al., 2020) using the Biomart tool (www.ensembl.org/biomart/), using human as reference. For each orthologous gene set, we downloaded exonic sequences for each species through Ensembl v83 (Table S5). For genes with multiple transcripts, we chose the longest coding sequence. We then used the *TBA* tool (Blanchette et al., 2004) to create multiple sequence alignments (MSA) for each orthologous gene set. Finally, we used in-house Python scripts to process Ensembl v83 GTFs of each species using the information from the MSA results, so as to include only exonic sequences that were fully alignable across all the species involved. Indels in the MSA (gaps inferred in one or multiple species) were removed. Finally, gene records resulting in sequences <100 bp were filtered out. These processed GTF's were used in the subsequent RNA-Seq data quantification steps.

***Constraints caused by the use of 1:1 orthologous genes:*** The reason behind relying on 1:1 orthologous genes was to readily identify and interpret convergent expression changes across species. Consequently, we excluded expression changes in lineage-specific genes and genes with paralogs. Such genes might be evolving rapidly and in testis-specific, cell-autonomous fashion (Brawand et al., 2011), and possibly under sexual selection. We note that, if the expression of genes with paralogs tends to evolve in more lineage-specific fashion than the expression of 1:1 orthologs, we could be overestimating the proportion of convergent patterns in testis transcriptome divergence. Also, the presence of distant mammalian species in some analyses (marmoset, gray short-tailed opossum and platypus), reduced the number of orthologous genes and the length of exons that are reported in the resulting final transcriptome annotations.

***Primate dataset no. 1:*** This RNA-Seq dataset was published in (Brawand et al., 2011). The testis subset includes 2 adult humans (*Homo sapiens*), 2 adult chimpanzees (1 *Pan paniscus* and 1 *Pan troglodytes*, which we do not distinguish as explained below), 1 adult gorilla (*Gorilla gorilla*), 2 adult rhesus macaques (*Macaca mulatta*). The prefrontal cortex subset includes 4 male humans, 6 male chimpanzees, and the same number of individuals for the other species as the testis subset. The data was produced on the Illumina Genome Analyser IIx platform. We downloaded the raw data for the brain and testis samples from NCBI GEO database (www.ncbi.nlm.nih.gov/geo/) with accession number GSE30352 raw reads from each individual were mapped to Ensembl v83 genomes and transcriptomes (see Table S5 for the genome assemblies used) using *TopHat2* (Kim et al., 2013) software, allowing maximum of 1 mismatch and 2 multi-hits for the reported alignments, limiting alignments to the annotated splice junctions ("–no-novel-juncs"), and changing the minimum and maximum intron length parameters for spliced reads to 40bp and 1Mb, respectively. Resulting alignments were further filtered to only include unique mappings using auxiliary fields in the BAM file output from the aligner. The filtered alignments were then quantified as FPKM expression on the gene level with *Cufflinks* (Trapnell et al., 2010), using the custom GTF file A for primates and also the custom GTF file B for all mammals included in the study. The resulting expression values were transformed as log2(x+1), where x stands for FPKM values, and quantile-normalized using the "preprocessCore" R package (Bolstad, 2019; Gautier et al., 2004).

***Primate dataset no. 2:*** This microarray dataset was published in (Khaitovich et al., 2005). It includes 6 adult human and 5 adult chimpanzee cerebral prefrontal cortex and testis tissue samples. The data was generated using Affymetrix Human HGU133Plus2 microarrays. We downloaded the raw CEL files from EBI ArrayExpress (www.ebi.ac.uk/arrayexpress/), under accession E-AFMX-11. We masked probes that do not match either human or chimpanzee genomes (GRCh38 and panTro2), and used a probeset definition based on Ensembl genes v83 (http://www.ensembl.org/), as described in (Somel et al., 2009). Affymetrix CEL files were processed using the Bioconductor "affy" package "rma" function (Gautier et al., 2004), which includes background subtraction, log transformation, normalization, and summary across probesets.

***Mouse, opossum and platypus testis:*** This RNA-sequencing dataset includes the same individuals described above in "primate dataset 1", as well as additional non-primate amniotes, and was published in (Brawand et al., 2011). We obtained the raw data from NCBI GEO database (www.ncbi.nlm.nih.gov/geo/) with accession number GSE30352. It was produced using bulk testis samples from adult individuals. Among the non-primate species, we used testis data from two adult house mice (*Mus musculus*), two gray short-tailed opossum (*Monodelphis domestica*) and three platypus (*Ornithorhynchus anatinus*) individuals. The data was generated as described above, and we followed the same steps used in preprocessing "primate dataset no. 1", the only difference being the use of custom GTF file B for all mammals in the quantification step.

***Marmoset testis:*** This RNA-Seq dataset was published in (Bellott et al. 2014) and was produced using bulk testis samples from a single adult common marmoset individual

(*Callithrix jacchus*). The data was generated on the Illumina MiSeq platform. The raw data was downloaded from NCBI Single Read Archive (http://www.ncbi.nlm.nih.gov/sra) with accession number SRX335333. The RNA-Seq data was processed as described above for the "primate dataset no. 1" using the custom GTF file created for marmoset.

***Rat testis:*** This dataset represents bulk testis samples from 2 adult rats. It was generated using Affymetrix Rat 230.2 microarrays and published in (Chalmel et al., 2007). Rat Affymetrix CEL files were downloaded from EBI ArrayExpress under accession number E-TABM-130. These were processed using the Bioconductor "affy" package "rma" function (Gautier et al., 2004). Affymetrix probeset IDs were mapped to rat genes and further to orthologous human Ensembl genes following the above-described procedure. We used the two adult rat bulk testis transcriptome profiles from this dataset.

***Mouse testis development:*** This microarray dataset was published in (Schultz et al., 2003). It includes 15 mice testis samples from individuals with ages ranging from 0 to 60 days. Affymetrix CEL files were downloaded from the NCBI GEO database (www.ncbi.nlm.nih.gov/geo/) with accession number GSE640, processed using the Bioconductor "affy" package "rma" function (Gautier et al., 2004). Affymetrix probeset IDs were mapped to Ensembl gene IDs (Ensembl v.83) through the Ensembl Biomart (www.ensembl.org/biomart/). For each gene that matched multiple probesets, we chose the single probeset with the highest mean expression level. We then mapped mouse genes to orthologous human genes, only including 1:1 orthologs as defined by Ensembl.

***Mouse testis cell types:*** We used two datasets published in (Chalmel et al., 2007) and (Namekawa et al., 2006). Both datasets include expression profiles from adult mouse spermatogonia, spermatocytes, and spermatids. In addition, the dataset published in (Chalmel et al., 2007) includes purified adult mouse Sertoli cells. Both studies had used Affymetrix Mouse 430.2 microarrays. Affymetrix CEL files were downloaded from EBI ArrayExpress under accession number E-TABM-130 and NCBI GEO under accession number GSE4193, and processed using the Bioconductor "affy" package "rma" function (Gautier et al., 2004). Affymetrix probeset IDs were mapped to mouse genes and further to orthologous human Ensembl genes following the above-described procedure. We jointly processed data from the two studies, as explained below.

***Cell-specific gene expression from human, macaque and mouse:*** This data was published in (Lesch et al., 2016). Raw data was acquired from the NCBI GEO database under accession number GSE68507. Except for one of the mouse samples, which was published before and sequenced on Illumina Genome Analyzer IIx platform, all libraries were sequenced on the HiSeq 2500 platform. The data included in this study were RNA-Seq profiles of pooled pachytene spermatocyte and round spermatid cells, from 3 humans, 2 rhesus macaques and 2 mice. The RNA-Seq data was processed as described above for the "primate dataset no. 1", the only difference being the use of custom GTF file B for all mammals in the quantification step.

***Human testis development:*** This data was published in (Cardoso-Moreira et al., 2019). All libraries were sequenced on the Illumina HiSeq 2500 platform. The samples used here included testis whole tissue RNA-Seq data from 14 individuals, between 221 postnatal days to 55 years of age. Raw data were downloaded from EBI ArrayExpress under accession number E-MTAB-6814, samples belonging to infants and older individuals were selected (i.e. prenatal samples excluded) and processed as described above for the "primate dataset no. 1".

### Combining datasets, species choice, and phylogenies

***Bonobo and chimpanzee:*** In "primate dataset no. 1" there existed a single bonobo, or pygmy chimpanzee (*Pan paniscus*). Studying this individual's bulk testis transcriptome profile relative to those of other primates, we observed that it fell within the variation of the common chimpanzee, *Pan troglodytes* (Figure S1). We therefore treated all *Pan* genus individuals as a single taxon and refer to this group as "chimpanzee" for simplicity.

***Combining primate testis datasets:*** Here our goal was to combine primate dataset no. 1 (RNA-Seq) with primate dataset no. 2 (microarray), such that we would retain biologically relevant variance while removing the effect of the platform differences, and consequently increase statistical power to study inter-species divergence. We merged the two datasets with a custom normalization approach, leveraging the fact that both datasets included adult human and chimpanzee testis samples.

1. We first filtered both datasets by removing genes that have 0 expression measured across all the individuals in that dataset. Note that the RNA-Seq dataset is already

filtered to include only 1:1 orthologs with alignable transcript sequence length ≥100 bp.

2. We joined the two datasets based on Ensembl genes common to both,. The resulting dataset contained 7,305 genes, with 8 humans, 7 chimpanzees, 1 gorilla and 2 rhesus macaques. This version will be referred to as “merged” in the following. We further refer to its components corresponding to primate dataset no. 1 and 2 as "subdatasets".
3. For each gene in the merged dataset, we calculated the mean expression level of all humans and chimps (n=15 in total) across all samples ($\mu_{h,c}$).
4. We then normalized each subdataset of the merged dataset using their human and chimpanzee subsets as reference, via the following steps:
    4.1. The subdataset corresponding to primate dataset no. 2 contained 6 humans and 5 chimpanzees; here we scaled each gene’s expression levels to mean=0 and sd=1.
    4.2. The subdataset corresponding to primate dataset no. 1 contained 2 humans and 2 chimpanzees, 1 gorilla and 2 rhesus macaques. Here we first scaled each gene’s expression levels to mean=0 and sd=1. We then subtracted the mean expression value for the 2 humans and 2 chimpanzees from all individuals’ expression values.

    The end result was that in both subdatasets, the human and chimpanzee individuals’ mean expression was 0.
5. We added to each gene’s expression levels the same gene’s $\mu_{h,c}$ value calculated at step (3). Thereby we rescaled the expression matrix to its original data range.

We analyzed overall similarities among the samples’ expression profiles in this dataset, both using principal component analysis and also using hierarchical clustering based on Euclidean distance, employing the “prcomp” and “hclust” functions, respectively, in the R base package “stats”. The results indicated that samples cluster according to their species identity and not dataset origin (Figure S1). The hierarchical clustering tree phylogeny also reflected the known species phylogeny.

In addition, we tested each gene in the combined primate testis dataset using two-way ANOVA including species and batch effects, employing the “aov” function in the R base package “stats”. Across all the 7,305 genes in the dataset, 4,081 (55.87%) showed a significant species effect (two-way ANOVA, at BH-corrected $q < 0.1$), while only 5 (0.07%) showed a significant batch effect. This indicated that our approach effectively removed major sources of between-dataset technical variance.

***Combining primate brain datasets:*** We applied the same procedure as above to combine the primate datasets no. 1 and 2 including the brain data. The combined dataset contained 7,212 genes, with 10 humans, 11 chimpanzees, 1 gorilla and 2 rhesus macaques.

***Combining mouse cell type datasets:*** Previous reports indicate that the main shift in transcriptome profiles during spermatogenesis occurs during meiosis – such that pre-meiotic and somatic cell types' profiles cluster together, while meiotic and post-meiotic cell types' profiles create a distinct cluster (Chalmel et al., 2007). Given this result, we combined the two mouse testis cell type datasets described above. Each dataset contained 4 pre-meiotic/somatic and 4 meiotic/post-meiotic samples – thus maintaining balance. To remove technical influence on gene expression, in each dataset we scaled each genes' mean to 0, and standard deviation to 1, and then merged the two matrices in a similar method as described above for combining primate bulk testis datasets by leveraging the fact that two datasets contained both spermatogonia and spermatid samples. The combined dataset included 8 PRE and 8 POST cell type expression profiles mapped to 14,934 human 1:1 orthologous Ensembl genes. Using hierarchical clustering as described above, we determined that PRE and POST samples cluster together (Figure S5), and dataset origins cannot be separated.

***Rationale for excluding marmoset, opossum and platypus from the main analyses:*** Although our study focuses on human, chimpanzee, gorilla, rhesus macaque (Old World anthropoids), we also analyzed testis transcriptome data from the house mouse, the common rat, the common marmoset, gray short-tailed opossum, and platypus. The house mouse and the common rat are known to mate highly promiscuously throughout the year (Kenagy and Trombulak, 1986; Chalmel et al., 2007). The common marmoset is generally assumed to be monogamous (Harcourt et al. 1981; Lukas and Clutton-Brock 2014), but mating competition between related males in the same group has been reported (de Sousa et al., 2009; Baker et al., 1999). Marmoset relative testis mass is reported to lie slightly below the mammalian average (Harcourt et al. 1981), but it's testis size also varies among males depending on breeding status (Araújo and De Sousa, 2008). The gray-tailed opossum is solitary; it has been noted to mate promiscuously in captivity (Jones et al., 2003), but has below-average relative testis mass (Rose et al., 1997). The platypus is also solitary. Although direct observations of mating behavior in the wild are not extensively reported, its large relative testis mass has been considered a sign of promiscuity (Rose et al., 1997; Ashwell, 2013). However, a study done on platypuses in captivity reported that a male platypus will try to mate with remaining unmated females present in the area whereas females will show avoidance after copulation therefore being unavailable for successive mating attempts from different males (Thomas et al., 2018). The fact that females become unavailable after they mate, points to polygny and thus, a single-male mating strategy. At the same time, the pronounced sexual dimorphism and the overlap between the period in which male platypuses have venom in their spurs and their mating season suggest physical competition with other males (Grant and Smith, 1998), but possibly not sperm competition.

When we compared bulk testis transcriptomes from these species to those of humans and chimpanzees, we found the two rodents showed significantly higher affinity to the chimpanzee, the platypus marginally significant affinity to the human, while the opossum and marmoset were equivocal (Figure S3). The result for the rodents is consistent with their multi-male mating strategies and high relative testis mass. Meanwhile, among the marmoset, opossum, and platypus, only the latter may follow expectations based on mating behavior and/or relative testis mass, assuming the platypus is polygynous (Thomas et al., 2018). There could be multiple possible explanations for the apparent inconsistencies: (a) the marmoset and opossum individuals might have been sampled within or outside of the breeding season, which might alter tissue composition and transcriptomes relative to the average testis state of the species (depending on the breeding season length and plasticity of testis anatomy and histology), (b) in the case of the marmoset, the sampled individual may have been a breeding male, known to have larger testicles than those of non-breeding males (Araújo and De Sousa, 2008), (c) the opossum may have been a juvenile, (d) the convergent patterns of gene expression evolution and cell type shifts observed for the Old World anthropoids and rodents may not apply to these three species. Without additional information regarding the exact characteristics of the individuals sampled, we are unable to evaluate these possibilities and interpret the observed inter-species transcriptome similarities. For these reasons we opted to exclude marmoset, opossum and platypus from further analyses.

***Biases due to confounding technical effects:*** The two-way ANOVA analysis described above indicated that technical biases were negligible in the combined primate dataset. Meanwhile, we did not attempt to combine other datasets directly (e.g. mouse cell type and primate testis). Here, instead, we compared data from outgroup species (e.g. mouse or marmoset) to the combined primate dataset, which contained a balanced number of humans and chimpanzees (note that human and chimpanzee numbers are balanced also within primate dataset 1 and within primate dataset 2). Thus, when comparing outgroup species (e.g. mouse or marmoset) to humans vs. chimpanzees, possible technical differences between the outgroup data and the primate data will not cause a bias towards either humans or chimpanzees.

***Phylogenies:*** For the convergence tests, we used the phylogenetic structures available at the TimeTree (timetree.org) database (Kumar et al., 2017) (Table S3).
